# Next generation SARM1 knockout and epitope tagged CRISPR-Cas9-generated isogenic mice reveal that SARM1 does not participate in regulating nuclear transcription, despite confirmation of protein expression in macrophages

**DOI:** 10.1101/2021.08.25.457655

**Authors:** Ciara G. Doran, Ryoichi Sugisawa, Michael Carty, Fiona Roche, Claire Fergus, Karsten Hokamp, Vincent P. Kelly, Andrew G Bowie

## Abstract

SARM1 is an ancient and highly conserved TIR-domain containing protein, with a diverse range of proposed roles in both innate immunity and neuronal death and degeneration. Murine SARM1 has been reported to regulate the transcription of specific chemokines in both neurons and macrophages, however the extent and mechanism by which SARM1 contributes to transcription regulation remains to be fully understood. Here, using RNA sequencing we identify differential gene expression in bone marrow-derived macrophages (BMDM) from C57BL/6 congenic 129 ES cell-derived *Sarm1*^-/-^ mice compared to wild type (WT). However, we show that passenger genes which are derived from the 129 donor strain of mice flank the *Sarm1* locus, confounding interpretation of results, since many of the identified differentially regulated genes come from the region containing passenger genes. To re-examine the transcriptional role of SARM1 in the absence of such passenger genes, we generated three different *Sarm1*^-/-^ mice using CRISPR/Cas9 technology. Vincristine treatment of *ex vivo* cultured post-natal neurons from these mice confirmed SARM1’s previously identified key function as an executor of axon degeneration. However, using these mice, we show that the absence of SARM1 has no impact on transcription of genes previously shown to be altered in macrophages or in the brainstem. To gain further insight into SARM1 function, we generated and characterized a mouse expressing epitope-tagged SARM1, as it has been difficult to date to confirm which cells and tissues express SARM1 protein. In these mice we see high SARM1 protein expression in the brain and brainstem, and lower but detectable levels in macrophages. Overall, the generation of these next generation SARM1 knockout and epitope-tagged mice has clarified that SARM1 is expressed in mouse macrophages but has no general role in transcriptional regulation in these cells, and has provided important new animal models to further explore SARM1 function.

## Introduction

Sterile alpha and HEAT/Armadillo motif containing protein (SARM1) is an ancient and highly conserved immune protein, with orthologues in lower organisms including *C. elegans*, and *D. melanogaster* as well as mammals [1]. It is the most recently identified member of the toll-like receptor (TLR) adaptor family of proteins, and comprises two sterile alpha motif (SAM) domains, armadillo repeats, and a toll/interleukin-1 receptor (TIR) domain [2]. The SAM domains mediate homotypic protein-protein interactions, and thereby facilitate SARM1 oligomerisation into an octamer in both its active and inactive states [3] [4]. The presence of a TIR domain predicts a role for SARM1 in innate immune signalling. However, in contrast to the other TLR adaptor proteins, SARM1 does not facilitate TLR signal transduction. In fact, human SARM1 has been shown to antagonize TLR signalling through both TIR domain-containing adaptor-inducing interferon-β (TRIF) [5] and Myeloid differentiation factor 88 (MyD88) [6] through TIR-TIR interactions [7]. As well as having an opposing role in signalling to other TLR adaptors, SARM1 is also distinguished from the family by the presence of NADase activity within its TIR domain [8]. Interestingly, phylogenetic analysis revealed that SARM1 TIR domain clusters more closely with bacterial TIRs than other animal TIR domain-containing proteins, suggesting it emerged in animals from bacteria *via* lateral gene transfer [9].

It has become clear in recent years that the functions of SARM1 extend beyond innate immunity. SARM1 is highly expressed in neurons, and hence many studies focus on the roles for SARM1 in cell death and axon degeneration in the nervous system. SARM1 was shown to mediate the death of mouse hippocampal neurons following glucose and oxygen deprivation [10], and murine SARM1 has also been shown to drive programmed cell death in sensory neurons downstream of oxidative stress [11]. In addition to regulating neuronal cell death, SARM1 has been identified as the central executor of axon degeneration in response to a range of triggers. Following axotomy, the distal portion of the transected axon disintegrates by an active process known as Wallerian degeneration. In mice lacking SARM1, severed axons are protected from this and exhibit remarkable long-term survival [12]. *Sarm1*^-/-^ mice also show attenuated surrogate markers of axon degeneration and reduced functional deficits after traumatic brain injury [13]. Additionally, in the absence of SARM1, mouse neurons are protected from the axon degeneration which is induced by the chemotherapeutic agents vincristine [14], paclitaxel [15], and cisplatin [16]. This axon degeneration limits the usefulness of these chemotherapies, as it results in peripheral neuropathy. In these circumstances, absence of SARM1 is advantageous, as it ameliorates the pathological outcomes associated with axon degeneration. There are situations however when SARM1-induced axon degeneration is beneficial. Recently, it was demonstrated that SARM1 is protective in a mouse model of ulcerative colitis [17]. SARM1-mediated axon degeneration in the enteric nervous system resulted in reduced secretion of norepinephrine, a neurotransmitter which drives inflammatory IL-17 secretion from T_H_17 cells and ILC3s causing local inflammation in the colon. In addition, in the context of rabies infection SARM1 mediates axonal degeneration and so diminishes the spread of the virus [18]. The role of SARM1 in axon degeneration can be accounted for by the intrinsic NADase activity in the TIR domain. This NADase activity becomes activated when the inhibitory lock between the TIR domain and auto-inhibitory ARM domain is disrupted [19], resulting in catastrophic NAD and ATP depletion within the axon which is temporally coupled with axon degeneration [8].

Independent from its role in cell death and axon degeneration, SARM1 is also known to regulate transcription in neurons under certain conditions. Following traumatic injury of neurons, SARM1 is required for the induction of CCL2, CCL7, CCL12 and CSF-1 [20]. Remarkably, the expression of these chemokines and cytokines could be induced by forced dimerization of the SARM1 TIR domain alone. In addition, altered transcription has been seen in the brain of SARM1-deficient mice in a number of neurotropic viruses. Following challenge with West Nile Virus (WNV), *Sarm1*^-/-^ mice displayed reduced TNF production in the brainstem, which resulted in enhanced susceptibility to the infection [21]. In response to Vesicular Stomatitis Virus (VSV), *Sarm1*^-/-^ mice showed reduced expression of a range of cytokines in the brain and, in contrast to WNV, this was protective and *Sarm1*^-/-^ mice did not succumb to infection [22]. We also previously described a role for SARM1 in regulating transcription in immune cells [23]. We reported that following stimulation of pattern recognition receptors with a range of ligands, *Sarm1*^-/-^ bone marrow-derived macrophages (BMDM) showed reduced *Ccl5* expression compared to WT cells, for both TLR-dependent and -independent stimulation of cells, suggesting a direct transcriptional role for SARM1 for select genes.

Compared to mechanistic insights on how SARM1 modulates axonal degradation that have arisen from elucidation of the structure and enzyme activity of SARM1 protein, it is unclear how SARM1 may regulate transcription, and how general or specific this regulation might be. To address this, we performed RNA sequencing on BMDM from TLR-stimulated WT and 129 ES cell-generated *Sarm1*^-/-^ mice backcrossed onto a C57BL/6 background (herein referred to as B6 congenic *Sarm1*^-/-^ mice). This revealed that a disproportionate fraction of differentially expressed genes (DEGs) flanked the *Sarm1* locus. A high density of single nucleotide polymorphisms (SNPs) and insertions and deletions (indels) associated with 129 mice were observed on either side of the *Sarm1* locus, indicating that these are passenger genes neighbouring the *Sarm1*^-/-^ mutation that were not removed during backcrossing to C57BL/6 mice. Three independent SARM1 deficient mice generated by CRISPR/Cas9-mediated genome editing confirmed that transcriptional phenotypes which we previously attributed to the absence of SARM1 were actually accounted for by the presence of passenger genes.

Further, there has been uncertainty as to where SARM1 is expressed, which has been an obstacle in gaining greater understanding of SARM1 function. The lack of a reliable commercially available antibody has precluded comprehensive investigation into which tissues and cells express SARM1. By generating a mouse expressing epitope-tagged SARM1 endogenously, we clarify here that SARM1 is indeed expressed in macrophages as well as the brain. We verify that macrophages from these mice exhibit normal cytokine induction following TLR stimulation, and show that the epitope tag does not interfere with SARM1 function in axon degeneration in primary neurons.

## RESULTS

### *Sarm1*^-/-^ macrophages show reduced expression of *Ccl5*, but not *Tnf* or *Il6*, compared to WT cells

We previously demonstrated that the expression of *Ccl5* is diminished in BMDM from *Sarm1*^-/-^ mice compared to WT mice following stimulation with a diverse range of both TLR and non-TLR ligands [23]. Here, we found that *Ccl5* transcription was significantly reduced in *Sarm1*^-/-^ pBMDM relative to WT following TLR7 stimulation with CL075 and TLR4 stimulation with MPLA over the course of 24 hours (Figures 1A and 1C). Similarly, following TLR4 stimulation with LPS *Ccl5* expression was reduced at the mRNA level, and while it fell short of statistical significance it is a consistent trend (Figure 1B). The diminished transcription led to substantially reduced secretion of CCL5 protein by *Sarm1*^-/-^ BMDM relative to WT following stimulation with LPS, MPLA, or CL075 (Figures 1J-L). This reduced CCL5 expression was not reflective of a global impairment in induction of inflammatory genes in BMDM from *Sarm1*^-/-^ mice since TNF (Figures 1D-F and 1M-O) and IL-6 (Figures 1G-I and 1P-R) mRNA and protein expression were not consistently lower in these cells relative to WT counterparts. This does not support an inhibitory role for murine SARM1 in TRIF signalling, contrary to human SARM1 [5].

**Figure 1.**
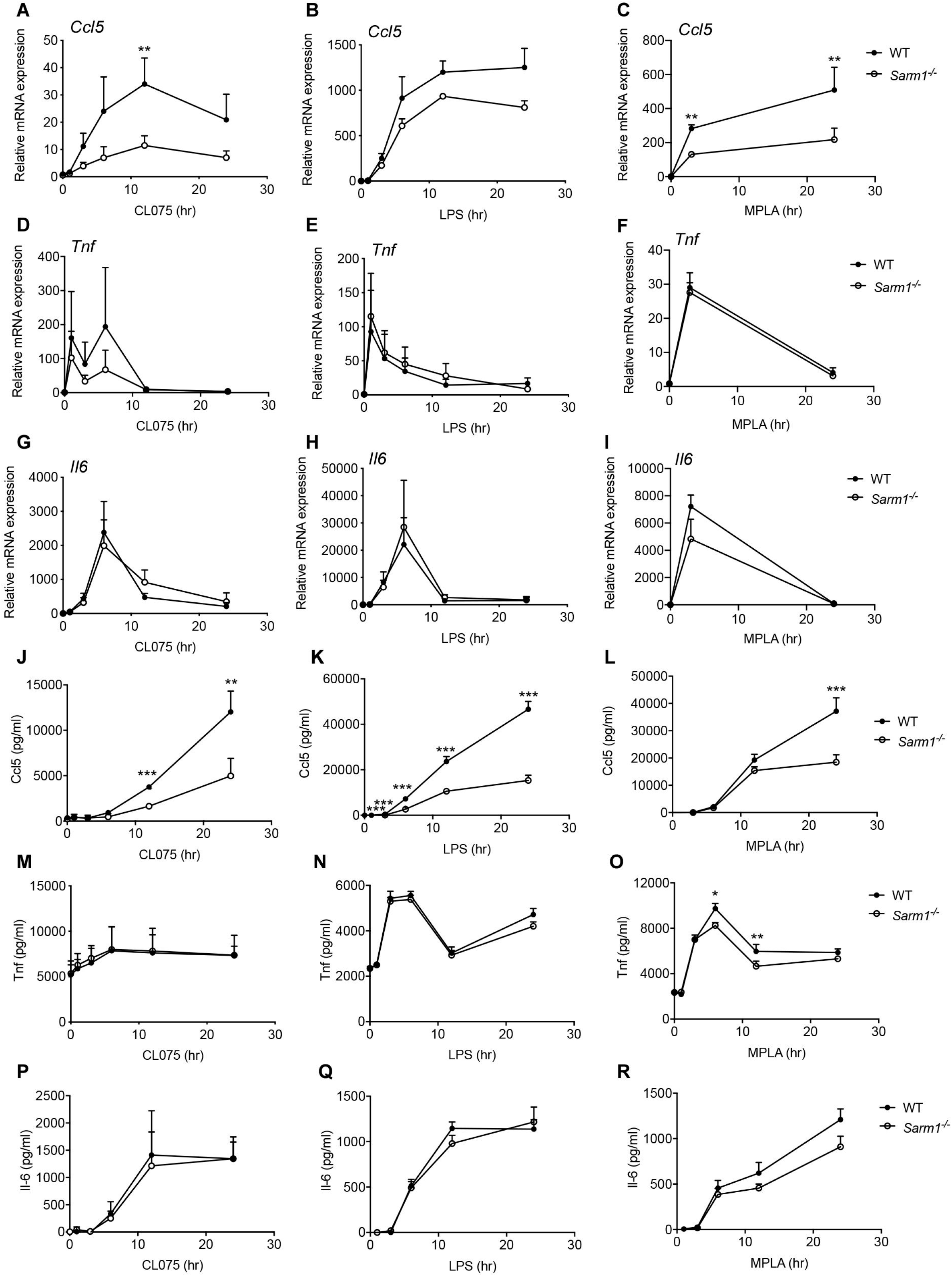
Ccl5 induction is reduced in BMDMs from *Sarm1*^-/-^ mice compared to WT mice. WT and *Sarm1*^-/-^ pBMDMs were stimulated with 5 μg/ml CL075 (A, D, G, J, M, P), 100 ng/ml LPS (B, E, H, K, N, Q), or 1 μg/ml MPLA (C, F, I, L, O, R) for the indicated times. *Ccl5* (A, B, C), *Tnf* (D, E, F), and *Il6* (G, H, I) mRNA were assayed by qRT-PCR, normalized to the housekeeping gene β-actin, and are presented relative to the untreated WT control. Supernatants were assayed for CCL5 (J, K, L), TNF (M, N, O), and IL-6 (P, Q, R) protein by ELISA. Graphs show mean ±SEM (n= 3-5) from at least 2 independent experiments. *p<0.05, **p<0.01, ***p<0.001 multiple Mann-Whitney tests; Holm-Šídák multiple comparisons. See also Fig S1.

We also previously showed that activation of transcription factors commonly associated with TLR-stimulated genes in macrophages, i.e NFκB and interferon regulators factors (IRFs), was normal in *Sarm1*^-/-^ BMDM. We therefore sought to determine the cellular localisation of SARM1 to understand how it may regulate transcription. While it has been reported that SARM1 can enter the nucleus to stabilize lamins [24], most reports show that SARM1 exists at the mitochondria [25, 26] and in the cytosol [27] . To assess if SARM1 could enter the nucleus to regulate transcription in BMDM, we used cellular fractionation of immortalized B6 congenic *Sarm1*^-/-^ BMDM stably overexpressing flag-tagged SARM1 to isolate a pure nuclear fraction and mixed cytoplasmic fraction. Flag-tagged SARM1 could not be detected in the nuclear fraction either before or after LPS treatment (Figure S1) showing that in BMDM, SARM1 does not reside at the nucleus, nor does it translocate there following LPS stimulation to regulate transcription.

### Passenger genes flank the *Sarm1* locus, and confound interpretation of differential gene expression

Knowing that TLR-stimulated CCL5 is diminished in B6 congenic *Sarm1*^-/-^ BMDM we wondered what other genes, if any, may be dependent on SARM1 for optimal expression, and the mechanism by which they could be SARM1-regulated. We therefore employed RNA sequencing to provide a broad, unbiased overview of transcription in BMDM from B6 congenic *Sarm1*^-/-^ mice. RNA from unstimulated BMDM was analyzed for differential gene expression to determine if any differences exist basally in the transcriptional landscape of BMDM from WT and B6 congenic *Sarm1*^-/-^ mice. Similarly, RNA from CL075-stimulated BMDM was analyzed for differential gene expression to identify potential SARM1-dependent TLR inducible genes. Differentially expressed genes (DEGs) from both comparisons were compiled into a single gene list, which was interrogated for patterns which may shed light on the mechanism by which SARM1 influenced their expression. No clear patterns emerged following transcription factor binding site analysis or pathway analysis, suggesting that these identified DEGs were not differentially expressed due to regulation of a known regulatory protein or transcription factor by SARM1. However, when a heatmap was generated with DEGs clustered by chromosomal location it became clear that chromosome 11, on which the *Sarm1* and *Ccl5* loci reside, was disproportionately represented (Figure 2A).

**Figure 2.**
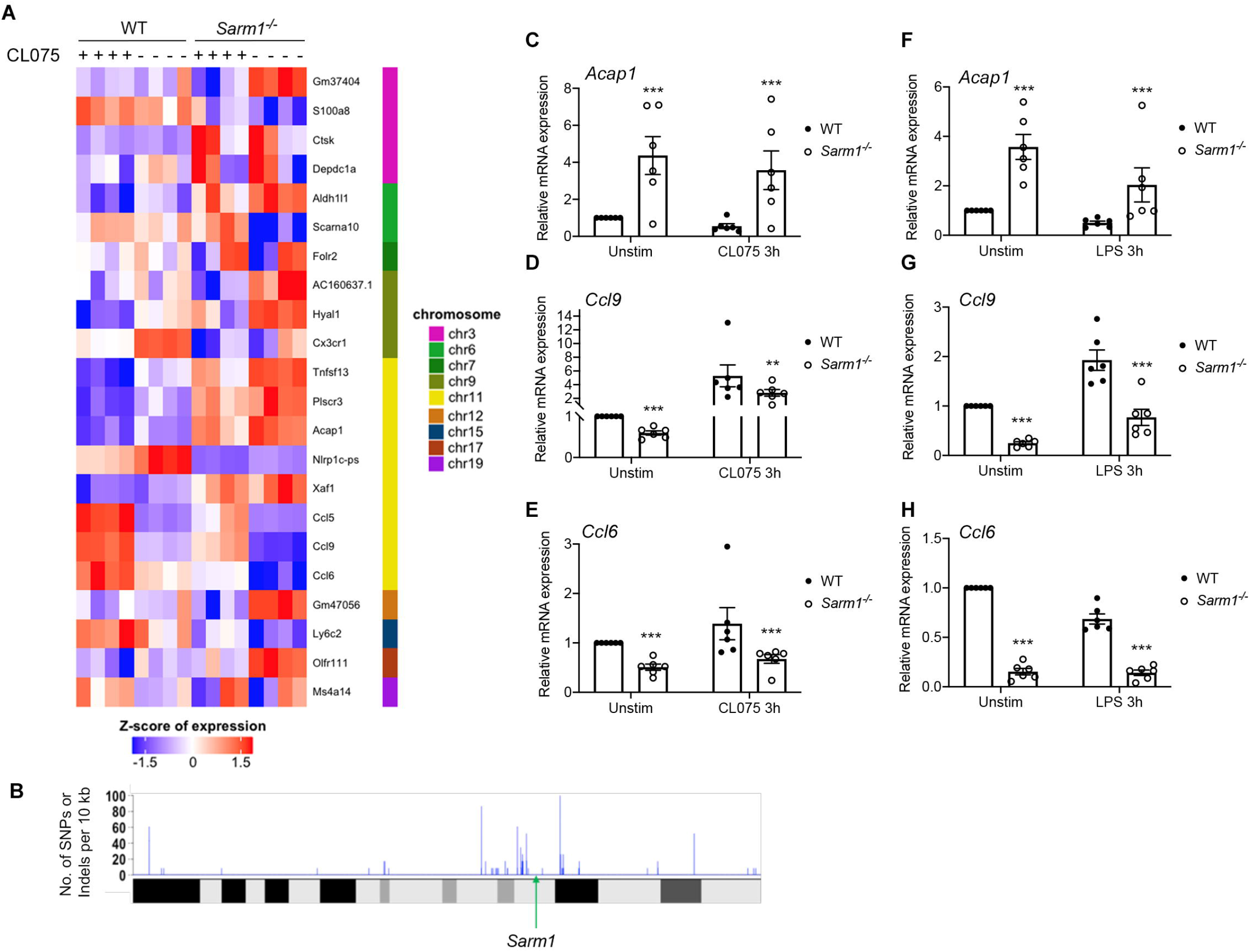
Transcriptome analysis of BMDMs from *Sarm1*^-/-^ mice compared to WT BMDMs reveals differentially expressed genes and passenger genes that cluster near the *Sarm1* locus. (A) Heatmap displaying genes which are differentially expressed between WT and *Sarm1*^-/-^ pBMDM, either basally or following CL075 stimulation for three hours, grouped according to chromosome. (B) Graph showing density of SNPs and indels per 10 kb section of chromosome 11 in *Sarm1*^-/-^ BMDM relative to the C57BL/6 reference sequence. The *Sarm1* locus is indicated by a green arrow. (C) WT and *Sarm1*^-/-^ pBMDMs were stimulated with 5 μg/ml CL075 (C, D, E) or 100 ng/ml LPS (F, G, H) for three hours, or medium as a control. *Acap1* (C, F), *Ccl9* (D, G), and *Ccl6* (E, H) mRNA were measured by qRT-PCR, normalized to the housekeeping gene β-actin, and are presented relative to the untreated WT control. Graphs show mean ± SEM (n= 6) from at least 2 independent experiments. **p<0.01, ***p<0.001, multiple Mann-Whitney tests; Holm-Šídák multiple comparisons. See also Fig S2 and Fig S3.

This prompted a re-examination of the B6 congenic *Sarm1*^-/-^ mouse, which was generated using targeted gene disruption. In this mouse, exons 3 through 6 of the *Sarm1* locus were disrupted by replacement with a targeting vector containing a neomycin-resistance cassette in embryonic stem cells derived from an unspecified 129/SvJ mouse, known as the donor [10]. These cells were implanted to a C57BL/6 blastocyst, the recipient, and following germline transmission and at least 15 generations of backcrossing, a SARM1-deficient mouse on the C57BL/6 background was achieved. However, it is now well established that mice generated in this way retain donor-derived genetic material, termed passenger genes or passenger mutations, in the region flanking the targeted gene [28]. The probability of a gene being of donor origin depends on the proximity to the targeted locus and the number of generations of backcrosses to C57BL/6 mice. To investigate the presence of these passenger genes in our mice, we compared the RNA sequence from B6 congenic *Sarm1*^-/-^ BMDM to the C57BL/6 mm10 reference sequence. Using a sliding window approach we examined the density of single nucleotide polymorphisms (SNPs) and insertions/deletions (indels) per 10Kb along chromosome 11 in the RNA from BMDM from B6 congenic *Sarm1*^-/-^ mice, relative to the C57BL/6 reference sequence, and found a high density of SNPs and indels in the region flanking the *Sarm1* locus (Figure 2B). This pattern was not seen on other chromosomes (Figure S2).

Among the differentially expressed genes in this region were *Ccl6, Ccl9*, and *Acap1*, which are ~4.32, ~4.31, and ~2.955 cM respectively from the *Sarm1* locus, as determined by the JAX Mouse Map Converter (based on [29] and [30]). Assuming 16 - 20 generations of backcrossing to C57BL/6, the probability of a passenger gene being retained within 5 cM from the targeted gene is 46.33 – 37.74% [31]. Thus, we examined the sequence of RNA from B6 congenic *Sarm1*^-/-^ BMDM for 129-associated variants at these loci. When the RNA sequence from B6 congenic *Sarm1*^-/-^ BMDM was examined using the Integrative Genomics Viewer (IGV) from the Broad Institute, deviations from the reference sequence were visible in the *Acap1* and *Ccl9* loci (Figure S3A and S3B). By manual comparison, these deviations were found to correspond to 129-associated SNPs as listed by the Wellcome Sanger Institute’s Mouse Genomes Project [32]. No SNPs or indels were visible in the *Ccl6* locus in B6 congenic *Sarm1*^-/-^ BMDM (Figure S3C), which corresponds with the lack of exonic 129-associated variations listed in the Mouse Genomes Project. DEG analysis showed that *Ccl6* and *Ccl9* expression were lower in B6 congenic *Sarm1*^-/-^ BMDM compared to their WT counterparts, and that *Acap1* was more highly expressed (Figure 2A). We validated these differences by quantitative RT-PCR: following either CL075 or LPS stimulation, *Sarm1*^-/-^ BMDM showed significantly less *Ccl6* and *Ccl9* expression, and significantly greater *Acap1* expression, than their WT counterparts (Figure 2C-2H). Importantly, these differences also existed in unstimulated cells, which could have implications for experimental models using these mice. Overall, RNA sequencing revealed that a number genes are differentially expressed in B6 congenic *Sarm1*^-/-^ BMDM compared to WT cells, but it is unclear whether this is due to the absence of SARM1 or the presence of passenger genes which flank the *Sarm1* locus in these cells.

### Generation and validation of three SARM1-deficient mice using CRISPR/Cas9

To study the effect of SARM1-deficiency on transcription in BMDM without the confounding factor of passenger genes, new *Sarm1*^-/-^ mice were generated by CRISPR/Cas9-mediated genome engineering. Figure 3A shows the CRISPR/Cas9 targeting strategy used, which generated three lines with indels of 2 bp, 5 bp and 34 bp resulting in premature stop codons. Each of the three novel lines contained deletions within the first exon of the *Sarm1* locus; *Sarm1^em1.1Tftc^* (2 bp deletion), *Sarm1^em1.2Tftc^* (34 bp deletion) and *Sarm1^em1.3Tftc^* (5 bp deletion) (Figure 3B). These deletions each interrupt the recognition site of the restriction enzyme BsaWI, which facilitated the use of this enzyme in genotyping (Figure 3B). Heterozygous breeding pairs were established to produce WT and SARM1-deficient littermate mice (Figure 3C). As SARM1 is highly expressed in neurons, we verified that SARM1 protein expression was lost in the brain of each line by Western blot (Figure 3D). SARM1 is difficult to reliably and specifically detect using commercially available antibodies, so we generated a polyclonal antibody by immunizing chicken with human SARM1 TIR domain recombinant protein. Indeed, SARM1 could be detected in the brains of WT mice, but not of their *Sarm1^em1.1Tftc^, Sarm1^em1.2Tftc^* and *Sarm1^em1.3Tftc^* littermates, nor in *Sarm1*^-/-^ brains (Figure 3D).

**Figure 3.**
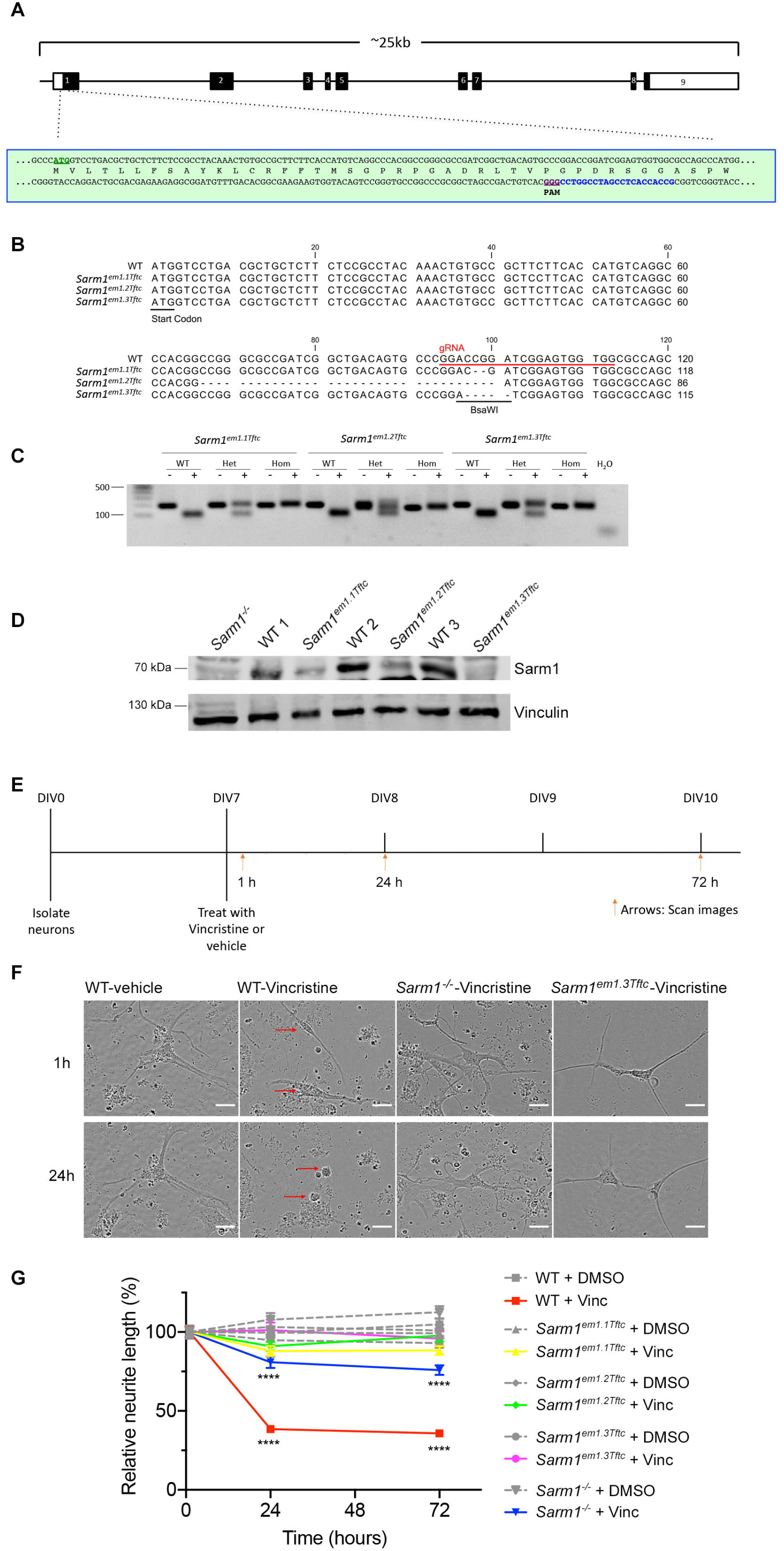
Generation of SARM1 knockout mice by CRISPR/Cas9 and validation of neuronal degeneration phenotype. (A) CRISPR/Cas9 targeting strategy for SARM1 knockout. The mouse *Sarm1* gene comprises 9 exons (numbered boxes). A gRNA target sequence was identified downstream of the ATG start site on the antisense strand (blue). CRISPR/Cas9 cutting at this site generated indels of 2 bp, 5 bp and 34 bp resulting in premature stop codons. (B) Multiple sequence alignment showing disrupted *Sarm1* locus compared to WT for *Sarm1^em1.1Tftc^* (2 bp deletion), *Sarm1^em1.2Tftc^* (34 bp deletion) and *Sarm1^em1.3Tftc^* (5 bp deletion). The start codon (black bar), the guide sequence (red bar) and the BsaWI, restriction enzyme site, used for genotyping are indicated. (C) Genotyping results for CRISPR littermates (WT, hets, homs). PCR amplicon from mouse genomic DNA was incubated with (+) or without (-) BsaWI enzyme. The amplicon containing WT sequence is recognised by the enzyme and shows a cleaved band whereas the disrupted sequence does not. (D) Immunoblot analysis of brain lysate for SARM1 and vinculin (loading control). WT1 is a littermate of *Sarm1^em1.1Tftc^*, same as WT2 and *Sarm1^em1.2TTtc^*, WT3 and *Sarm1^em1.3Tftc^* respectively. (E) Schematic of axon degeneration protocol. DMSO is used as a vehicle. Primary neurons DIV (days in vitro) 7 were treated with vincristine or vehicle and images are scanned after 1h, 24h and 72h for analysis used in (F-G). (F) Representative images of neurons after 1h or 24h treatment of vincristine or DMSO. Red arrows indicate cell bodies of WT neurons that after 24h vincristine treatment lost neurites. White scale bar, 25 μm. See also Movies S1 and S2. (G) Graph of relative neurite length over time for different mice and treatments. Neurite lengths are normalized to its mean of own 1h length as 100 (%). All data are mean ± SEM (n= 4~5). Data were tested with a 2-way ANOVA showing significant main effects of group F (9, 4973) = 59.02, P<0.0001; time F (2, 4973) = 25.08, P<0.0001; and interaction F (18, 4973) = 14.90, P<0.0001; Tukey’s multiple comparisons test, ****P < 0.0001 vincristine versus vehicle (DMSO) in WT and *Sarm1*^-/-^ at 24h and 72h. See also Figure S4 and Supplemental movies S1 and S2.

The role of SARM1 as a driver of axon degeneration has been extensively studied in the context of traumatic brain injury, axotomy, and mitochondrial dysfunction, but mainly in B6 congenic *Sarm1*^-/-^ mice. The chemotherapeutic drug vincristine is widely reported to induce SARM1-dependent axon degeneration [14, 33, 34]. We exploited this to assess and confirm the loss of SARM1-dependent axonal degeneration in the *Sarm1^em1.1Tftc^, Sarm1^em1.2Tftc^*, and *Sarm1^em1.3Tftc^* mice. Using live-cell imaging microscopy, we examined vincristine-induced neurite degeneration in post-natal primary neurons cultured for 7 days in vitro (DIV 7) from WT, B6 congenic *Sarm1*^-/-^ mice, and each of the three CRISPR knockout lines (Figure 3E). Neurons were treated with vincristine and scanned every six hours for three days, starting 1 hour post-treatment. WT neurons exhibited altered morphology and clear neurite loss in the 24 hours following vincristine treatment (Figure 3F, Supplemental movies S1 and S2). In line with the literature, *Sarm1*^-/-^ neurons were protected from axon degeneration following insult with the microtubule poison vincristine. Neurons from *Sarm1^em1.3Tftc^* mice were also robustly protected from axon degeneration (Figure 3F). In contrast to prenatal dorsal root ganglion neurons which are often used in such assays, fragments from degenerated primary neuron neurites disappear and couldn’t be counted to quantitate. Therefore, to quantitatively appraise the axon degeneration, we measured neurite length, which is commonly used to determine the effects of a particular substrate on neural outgrowth integrity. We measured neurite length at 1h, 24h, and 72h after vincristine treatment. Vincristine treatment resulted in dramatic and significant loss of neurite length in WT neurons after 24 hours which was maintained after 72 hours (Figure 3G; unnormalized and raw data for this is shown in Figure S4A and S4B). As expected from previous studies, *Sarm1*^-/-^ neurons were protected from loss of neurite length following vincristine treatment. Importantly, neurons from each of the three new CRISPR knockout lines retained their full neurite length, even 72 hours after vincristine treatment. Thus this assay demonstrated that neurons from *Sarm1^em1.1Tftc^, Sarm1^em1.2Tftc^*, and *Sarm1^em1.3Tftc^* mice faithfully recapitulate the axoprotective phenotype of *Sarm1*^-/-^ neurons.

### In the absence of passenger genes, SARM1-deficient BMDM display normal transcription

Having verified that SARM1 protein is not expressed in *Sarm1^em1.1Tftc^*, *Sarm1^em1.2Tftc^*, and *Sarm1^em1.3Tftc^* and having shown that the CRISPR knockout lines are phenotypically similar to *Sarm1*^-/-^ mice with regard to axon degeneration, we then sought to interrogate the transcriptional phenotype in BMDM without the confounding factor of passenger genes. In particular, since *Ccl5* resides on chromosome 11, less than 5 cM from the *Sarm1* locus, passenger genes could explain the differential *Ccl5* expression seen in *Sarm1*^-/-^ BMDM (Figure 1). Indeed, contrary to what was seen in *Sarm1*^-/-^ BMDM, *Ccl5* transcription and subsequent secretion was not significantly different in BMDM from *Sarm1^em1.3Tftc^* mice compared to their WT littermates following stimulation with LPS over a 24 hour period (Figures 4A and 4C). A similar phenotype was observed in BMDM from both *Sarm1^em1.1Tftc^* and *Sarm1^em1.2Tftc^* mice (Figure S5). The reduced *Ccl5* expression previously observed in B6 congenic *Sarm1*^-/-^ BMDM was therefore not as a result of SARM1-deficiency, but of passenger genes. We then measured expression of three genes which were identified by RNA sequencing as differentially expressed in *Sarm1*^-/-^ and validated by RT-PCR: *Acap1, Ccl9*, and *Ccl6*. Compellingly, all three genes were expressed equally in BMDM from *Sarm1^em1.3Tftc^* mice and their WT littermates, both basally and following LPS stimulation over a 24 hour time course (Figure 4E-G). Therefore, the previously observed differential expression resulted from the presence of passenger genes and not the absence of SARM1.

**Figure 4.**
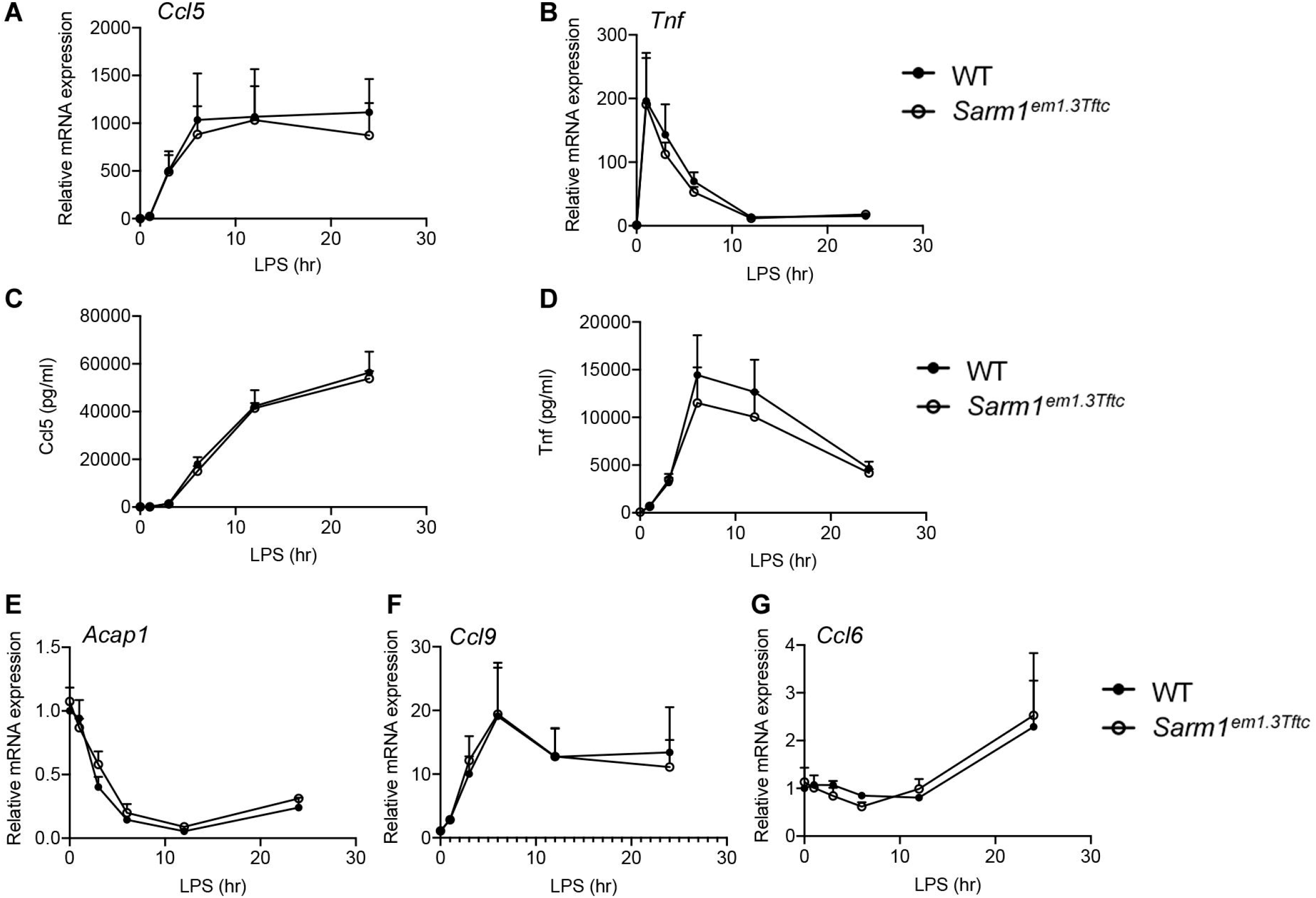
Macrophages from *Sarm1^em1.3Tftc^* mice show no defect in induction of select genes on chromosome 11 compared to WT littermates. (A-G) WT and *Sarm1^em1.3Tftc^* pBMDM were stimulated with 100 ng/ml LPS for the indicated times, or medium as a control. Expression of *Ccl5* (A), *Tnf* (B), *Acapl* (E), *Ccl9* (F), and *Ccl6* (G) mRNA in were assayed by quantitative RT-PCR, normalized to the housekeeping gene β-actin, and are presented relative to the untreated WT control. Supernatants were assayed for CCL5 (C) and TNF (D) protein by ELISA. Graphs show mean ±SEM (n= 3) from two independent experiments. No significant differences determined by multiple Mann-Whitney tests; Holm-Šídák multiple comparisons.

### Transcription is normal in the brainstem of *Sarm1^em1.1Tftc^, Sarm1^em1.2Tftc^*, and *Sarm1^em1.3Tftc^ mice*

Although SARM1 deficiency does not alter transcription in BMDM in the absence of passenger genes we wondered if differences in transcription would persist in the brain, where SARM1 is reported to be more highly expressed. Other transcriptional phenotypes previously ascribed to SARM1-deficiency in the brain have also been shown to be attributable to passenger genes in *Sarm1*^-/-^ mice. Recently, the García-Sastre lab generated two independent SARM1-deficient mice using CRISPR technology, and demonstrated that their previously published phenotype of altered cytokine and chemokine production in the brain of *Sarm1*^-/-^ mice following VSV infection [22] was not recapitulated where passenger genes are absent [35]. In that study, they found using RNA sequencing that a number of genes, including *Tfrc, Rps29, Rp138, Ndufb3*, and *Atp5k* were homeostatically differentially expressed in the brainstem of SARM1-deficient mice generated by CRISPR relative to WT. Having verified that *Sarm1^em1.1Tftc^, Sarm1^em1.2Tftc^*, and *Sarm1^em1.3Tftc^* mice do not express SARM1 protein in the brain, we therefore examined expression of *Ccl5, Tfrc, Rps29, Rpl38, Ndufb3*, and *Atp5k* in the brainstem in each of these independent lines. Using RT-PCR, we found no significant differences in expression of any of these genes in the brainstem of *Sarm1^em1.1Tftc^* mice compared to their WT littermates (Figure 5A-F), nor in the brainstems of *Sarm1^em1.2Ttc^* and *Sarm1^em1.3Tftc^* mice compared to their WT littermates (Figure S6). Thus data from our three independent SARM1-deficient mice do not support a role for SARM1 in transcription in the absence of passenger genes, neither in BMDM nor in the brainstem.

**Figure 5. Title.**
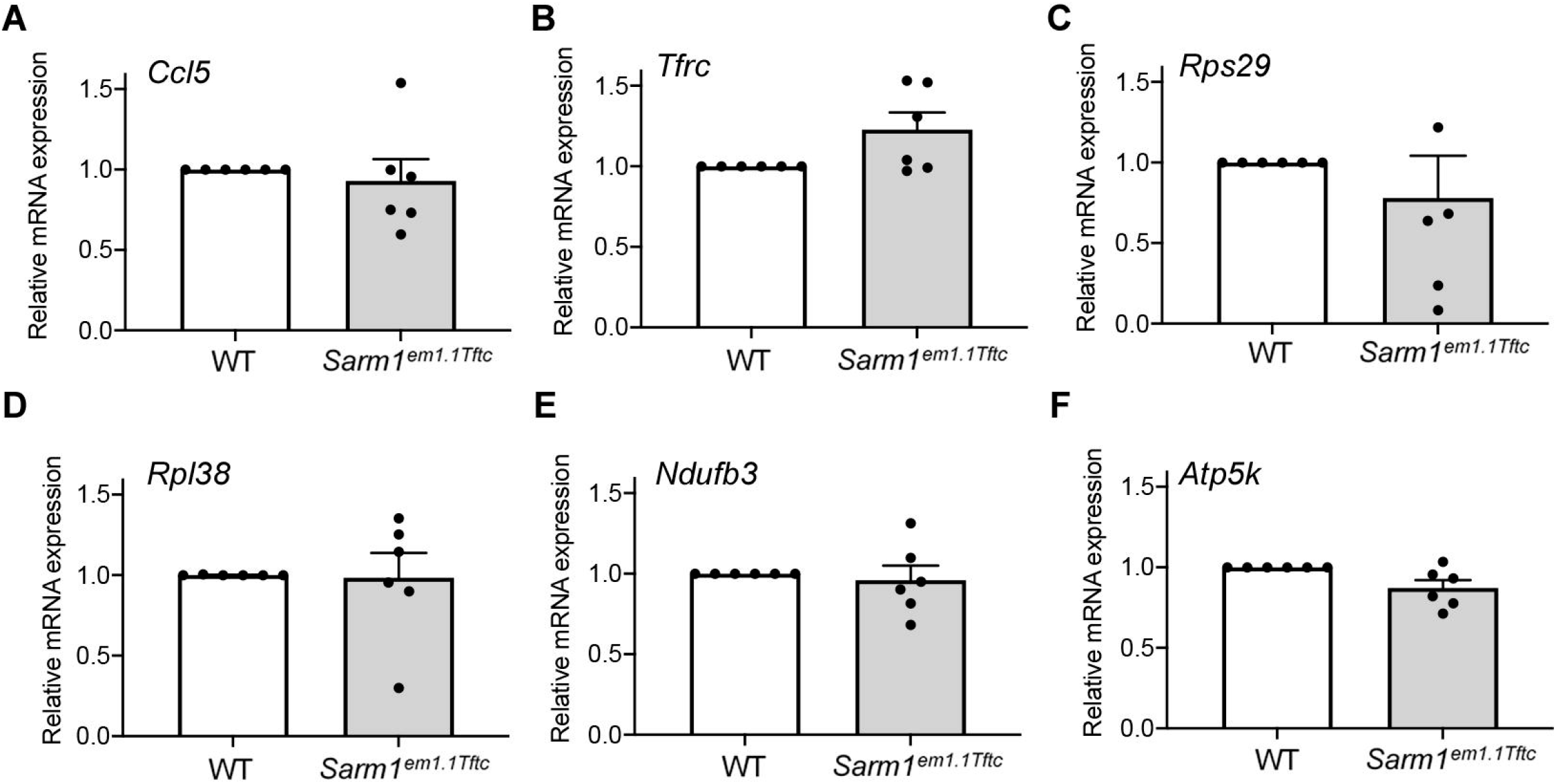
The absence of SARM1 does not result in altered transcription of select genes in the brainstem. (A-F) Expression of *Ccl5* (A), *Tfrc* (B), *Rps29* (C), *Rpl38* (D), *Ndufb3* (E), and *Atp5k* (F) in the brainstem of WT and *Sarm1^em1.1Tftc^* littermate mice were measured by qRT-PCR, normalized to β-actin, and are presented relative to the littermate WT. Data are mean ±SEM (n= 6), no significant differences determined by multiple Wilcoxon tests; Holm-Šídák multiple comparisons. See also Fig S6.

### Generation and validation of mouse expressing epitope-tagged SARM1

Our data from these three new CRISPR/Cas9-generated *Sarm1*^-/-^ mice confirms a role for SARM1 in neuronal axonal degeneration but does not reveal any phenotypic transcriptional differences between *Sarm1*^-/-^ and WT littermate BMDMs. Although we and others have reported roles for SARM1 in macrophages, the lack of a commercially available antibody which can reliably and specifically detect mouse SARM1 in tissues outside of the brain has hindered studies into the functions for SARM1 outside of the nervous system, including in macrophages. To overcome this, we used CRISPR technology to successfully generate a mouse expressing an epitope-tagged SARM1 endogenously, with a triple flag tag and double strep tag on the C-terminal end (Figure 6A), herein referred to as *Sarm1^Flag^* mice (see Methods for details on the targeting and repair strategy used). These *Sarm1^Flag^* mice showed similar *Sarm1* mRNA expression to their WT counterparts across a number of tissues, being most highly expressed in the brain, and expressed at a lower level in the liver, spleen and kidneys (Figure 6B).

**Figure 6.**
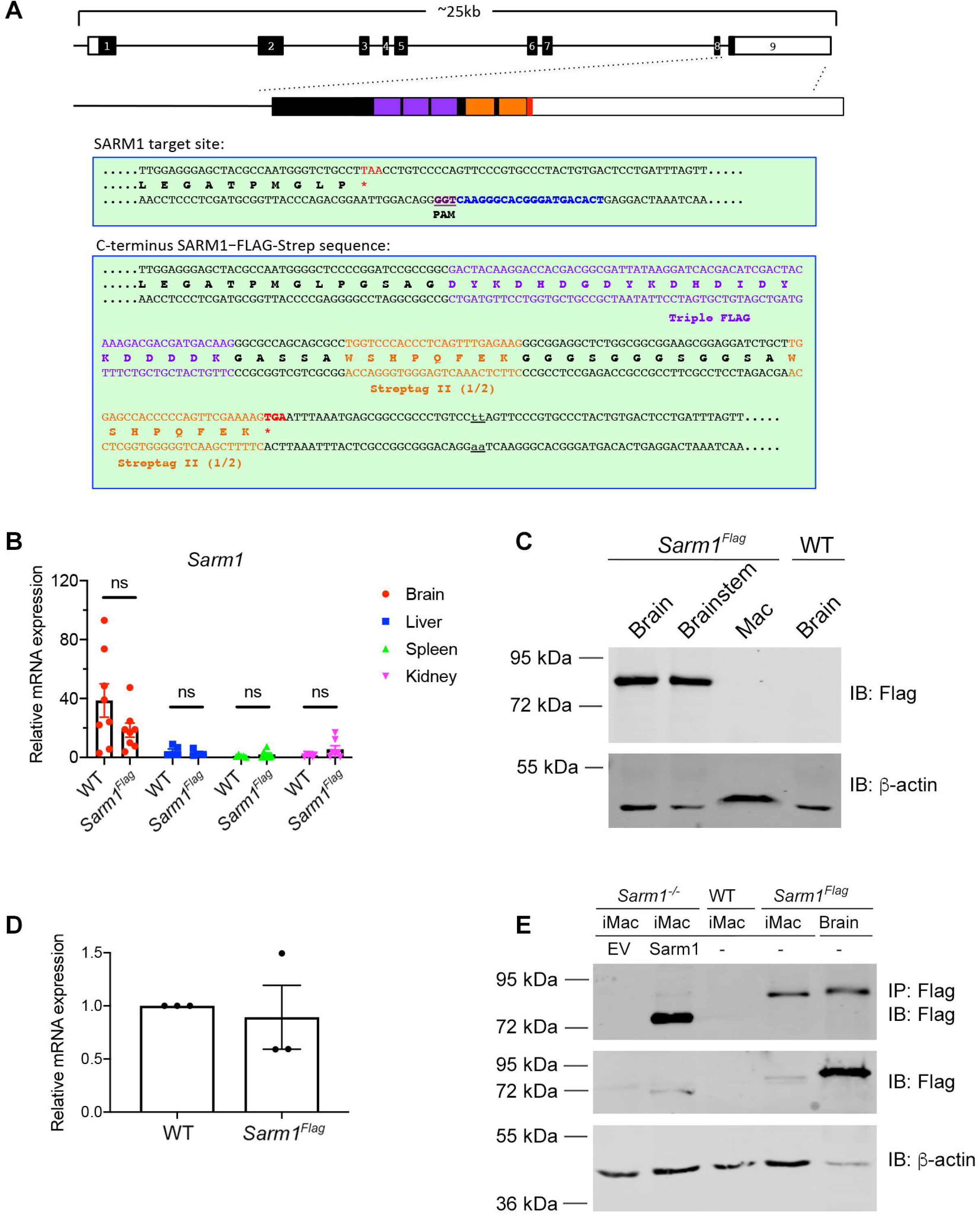
Generation of epitope-tagged SARM1 mice, *Sarm1^Flag^*. (A) CRISPR/Cas9 targeting strategy to generate SARM1 FLAG-Strep (*Sarm1^Flag^*) mice. The mouse *Sarm1* gene comprises 9 exons (numbered boxes). A gRNA target sequence was identified in exon 9 (blue) proceeding the stop codon. A repair template was used to insert a triple FLAG, double Streptag II (purple and orange boxes, respectively) at the C-terminus *via* homology directed repair. A cc > tt base change was engineered into the repair construct, underlined, to ablate the PAM sequence of the original gRNA target site. (B) Expression of *Sarm1* mRNA in brain, liver, kidney, spleen assayed by quantitative RT-PCR, normalized to the housekeeping gene β-actin, and are presented relative to the WT spleen. A two-tailed Mann-Whitney test was used to calculate the p values. ns, not significant. (C) Immunoblot comparing expression of flag-tagged SARM1 in brain, brainstem, and BMDM from *Sarm1^Flag^* mice, with WT brain lysate acting as a negative control. 30 μg of protein was loaded per sample as determined by BCA assay. β-actin was used as a loading control. Blot is representative of four independent experiments. (D) Expression of *Sarm1* mRNA in *Sarm1^Flag^* BMDM as measured by qRT-PCR, normalized to the housekeeping gene β-actin, and presented relative to the WT control. Data are mean ±SEM (n= 3) from two independent experiments (E) SARM1 is expressed weakly in BMDM compared to brain. Flag-tagged SARM1 was detected by immunoprecipitation in *Sarm1*^-/-^ iBMDM stably overexpressing SARM1 or *Sarm1^Flag^* iBMDM (iMac), and *Sarm1^Flag^* brain. β-actin was used as a loading control for input. Representative of 3 independent experiments.

We used the *Sarm1^Flag^* mice to interrogate SARM1 protein expression in the brain, the brainstem and BMDM. Western blotting showed that SARM1 is as highly expressed in the brainstem as it is in the cerebrum (Figure 6C). SARM1 was undetectable by anti-flag immunoblot in BMDM from *Sarm1^Flag^* mice (Figure 6C), although it was detectable at the mRNA level (Figure 6D). To overcome experimental constraints associated with limited primary BMDM numbers, we established immortalized BMDMs from *Sarm1^Flag^* mice and littermate WT mice. SARM1 was clearly detected in these immortalized *Sarm1^Flag^* BMDM using flag immunoprecipitation followed by anti-flag immunoblot, albeit at lower levels than in the brain of *Sarm1^Flag^* mice (Figure 6E). Hence, we confirmed SARM1 protein expression is not limited to brain.

To facilitate future studies using the *Sarm1^Flag^* mice, we performed experiments to confirm that SARM1-flag protein retained functionality in these mice. Firstly we assessed cytokine induction in pBMDM from *Sarm1^Flag^* mice following LPS stimulation and found that *Sarm1^Flag^* BMDM showed normal *Ccl5* and *Tnf* transcription and secretion (Figure 7A-D). Importantly, we used the vincristine-induced neurite degeneration assay which previously confirmed that this process is SARM1-dependent (Figure 3G) to assess whether SARM1-flag protein in the *Sarm1^Flag^* mice retained the ability to mediate axonal degeneration. We found that neurons from *Sarm1^Flag^* mice showed similar loss of neurite outgrowth to WT neurons after 24 and 72 hours, indicating that epitope-tagged SARM1 protein retains its ability to be activated and mediate axon degeneration (Figure 7E, unnormalized and raw data for this is shown in Figure S7). Thus, using these *Sarm1^Flag^* mice we clarify that SARM1 protein expression is not limited to the brain and show that the epitope tags allow for detection of SARM1 without interfering with SARM1 expression or neuronal functional. These mice therefore are a useful tool for further studies into SARM1 expression and function.

**Figure 7.**
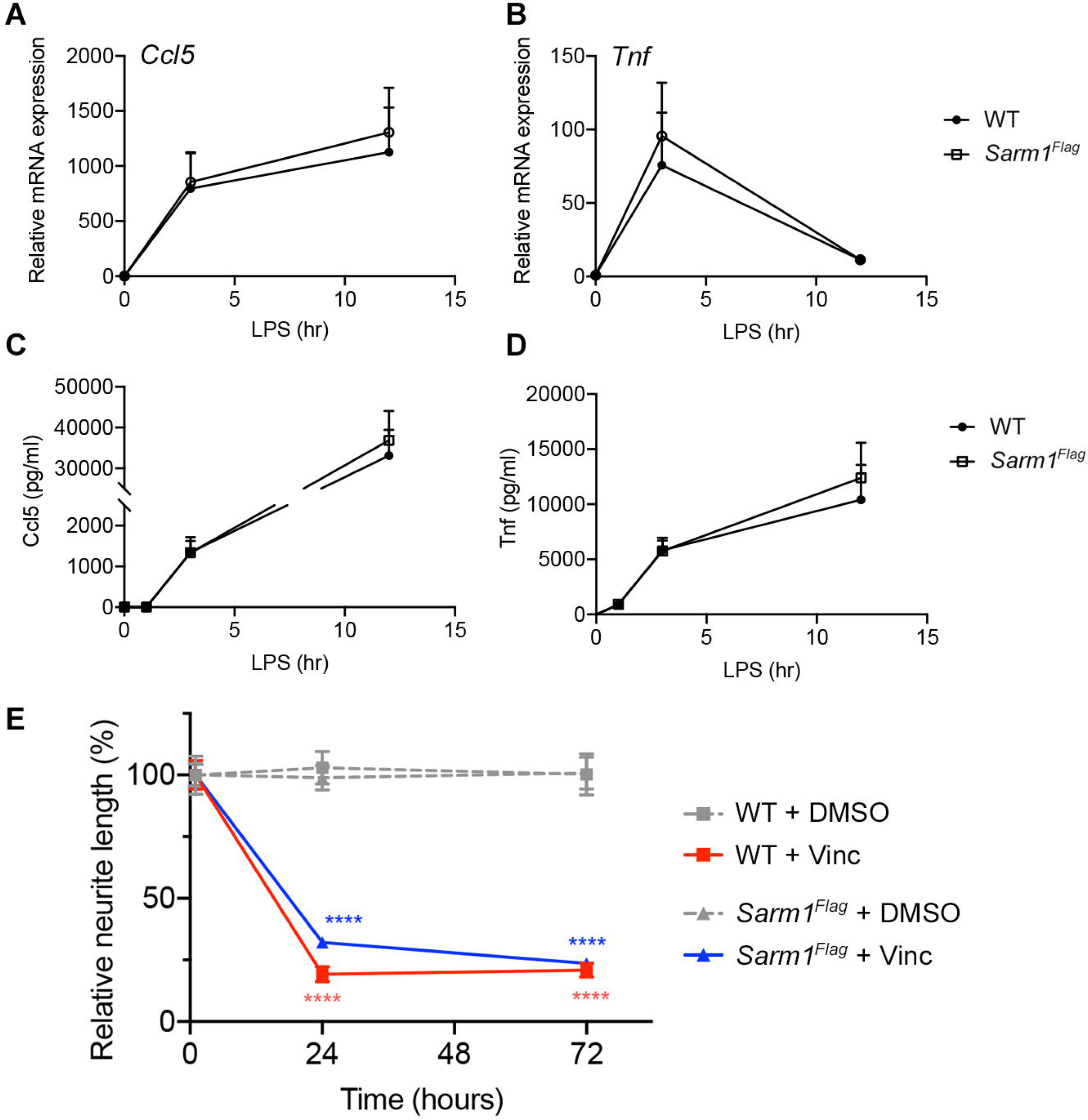
*Sarm1^Flag^* mice display similar gene induction and neuronal degeneration phenotypes to WT mice. (A-D) WT and *Sarm1^Flag^* BMDM were stimulated with 100 ng/ml LPS as indicated. Expression of *Ccl5*(A) and *Tnf* (B) mRNA were assayed by qRT-PCR, normalized to the housekeeping gene β-actin, and are presented relative to the untreated WT control. Supernatants were assayed for CCL5 (C) and TNF (D) protein by ELISA. Data are mean ±SEM (n= 3-4) from 2-3 independent experiments. No significant differences determined by multiple Mann-Whitney tests; Holm-Šídák multiple comparisons. (E) Graph of relative neurite length over time for different mice and treatments. Neurite lengths are normalized to its mean of own 1h length as 100%. All data are mean ± SEM (n= 2~4). Data were tested with a 2-way ANOVA showing significant main effects of group F (3, 920) = 82.79, P<0.0001; time F (2, 920) = 54.11, P<0.0001 and interaction F (6, 920) = 22.26, P<0.0001; Tukey’s multiple comparisons test, ****P < 0.0001 vincristine versus vehicle (DMSO) in WT and *Sarm1^Flag^* at 24h and 72h. See also Figure S6.

## DISCUSSION

We initially set out to understand the mechanism by which SARM1 positively regulates TLR-dependent *Ccl5* transcription in murine macrophages. Given that we previously demonstrated that transcription factor translocation to the nucleus was normal in *Sarm1*^-/-^ BMDM, but transcription factor and RNA polymerase II recruitment to the *Ccl5* promoter was diminished relative to WT [23] we had considered the possibility that SARM1 could reside at or enter the nucleus of BMDM following stimulation. There, we speculated that it may physically interact with transcription factors, or catalyze the local depletion of NAD, reducing its availability to enzymes which modulate chromatin accessibility, and thereby regulate transcription. However we found that SARM1 does not reside at the nucleus or translocate there following stimulation, so we therefore undertook a full analysis of the TLR-modulated transcriptome in WT and *Sarm1*^-/-^ BMDM in order to obtain insights into the role of SARM1 in regulating nuclear transcription. Through analysis of RNA sequencing data, it became apparent that proximity to the *Sarm1* locus was a common factor among a number of DEGs. Interrogation of the RNA sequence from *Sarm1*^-/-^ BMDM revealed a high density of SNPs and indels relative to the C57BL/6 reference sequence on chromosome 11, flanking the *Sarm1* locus. Further analysis revealed that a number of the DEGs had sequence deviations which corresponded to 129-associated variants, indicating that these were passenger genes and were introduced to the genome during the generation of the B6 congenic *Sarm1*^-/-^ mouse. Some DEGs did not display any sequence deviations, such as *Ccl6*, however their altered expression may be a result of SNPs or indels within promoters or enhancers, or disruption of the 3D chromosomal landscape. These differences would not be detectable by examination of the transcribed RNA alone. A number of genes were differentially expressed basally between BMDMs from WT and B6 congenic *Sarm1*^-/-^ mice, and this could have implications for the interpretation of experiments involving B6 congenic *Sarm1*^-/-^ mice or tissues or cells derived from these mice.

It has been well-established that generation of C57BL/6 congenic mice following target gene disruption in 129 ES cells results in residual donor genetic material in the recipient mouse in the form of passenger genes, and that the probability of a certain locus retaining this donor material is a function of its proximity to the target gene and the number of generations of backcrosses to the recipient mouse strain. In this case, a number of genes we observed to be differentially expressed in *Sarm1* BMDM by RNA sequencing lie near the *Sarm1* locus. *Ccl6, Ccl9, Acap1*, and *Ccl5* are all within 5 cM of the *Sarm1* locus, where the probability of genes being of 129 origin is ~46% [31], assuming 16 backcrosses to the C57BL/6 genome. The generation of three independent SARM1-deficient mice using CRISPR/Cas9 allowed us to delineate whether it was the absence of SARM1 or presence of passenger genes which accounted for differential expression. Our data from these mice did not support a role for SARM1 in transcription in BMDM. This is in agreement with the García-Sastre lab’s report that *Ccl3, Ccl4*, and *Ccl5* transcription are defective in B6 congenic *Sarm1*^-/-^ mice, but normal in CRISPR knockout mice [35].

Also in line with findings from the García-Sastre lab, we noted that an alternative *Xaf1* transcript with an extended 3’ UTR was expressed in B6 congenic *Sarm1*^-/-^ BMDM (data not shown). This has implications for a previous report that used these mice to implicate SARM1 in regulation of XAF1 and hence neuronal apoptosis [36]. In contrast to findings from the García-Sastre study [35], we did not observe any alterations in basal expression of brainstem *Tfrc, Rps29, Rpl38, Ndufb3* or *Atp5k* between WT and CRISPR-generated SARM1-deficient mice, across three independent lines. These differences between the two studies remain to be resolved. Our findings add to a growing body of literature implicating passenger genes as a possible cause of phenotypes previously attributed to the targeted gene in congenic mice [37–40].

While transcriptional phenotypes previously reported in *Sarm1*^-/-^ mice did not persist in our CRISPR-generated SARM1-deficient mice, the axoprotective phenotype did. We found that primary neurons from each of the three CRISPR-generated SARM1-deficient mice were robustly protected from vincristine-induced axon degeneration, which is the first demonstration of SARM1-dependent protection from chemotherapeutic agent-induced axonal loss outside of congenic *Sarm1*^-/-^ mice. As such, this demonstrates that these mice are a valuable tool which may be useful for further studies pertaining to murine SARM1 function in neurons, particularly where altered transcription due to passenger genes may confound interpretation results. Having such reliable animal models to probe SARM1 function *in vivo* is especially important and urgent currently since new data on SARM1 structure and function is rapidly emerging [41–44], leading to a heightened focus on SARM1 as a potential therapeutic target for neurodegenerative disease, traumatic brain injury and peripheral neuropathies [45, 46]

Comprehensive investigation into which cells and tissues mouse SARM1 is expressed in, particularly in regard to macrophages, has been hindered by difficulty in detecting SARM1 using commercially available antibodies, and has been also noted by others [35]. To overcome this, we generated to our knowledge the first epitope-tagged SARM1 expressing mouse using CRISPR/Cas9-mediated genome engineering. In this *Sarm1^flag^* mouse, we found robust SARM1 protein expression in the brainstem, where no transcriptional phenotype was observed, and in the brain. In BMDM where SARM1 expression has been contested, we saw that SARM1 was expressed, though to a lesser degree than in the brain. It was readily detectable by Western blot following immunoprecipitation in immortalized BMDM from *Sarm1^flag^* mice. We were unable to detect tagged SARM1 similarly in primary BMDM however, so it may be that immortalization increases the expression of SARM1. We verified that these mice resemble WT in both *Sarm1* expression and function. In the *Sarm1^flag^* mice, the mRNA levels of *Sarm1* were normal in brains, liver, spleen, and kidney. Further, we found that BMDM from *Sarm1^flag^* and WT mice showed similar cytokine induction following stimulation. Importantly, by characterizing axon degeneration in *Sarm1^Flag^* primary neurons, we could demonstrate that epitope-tagged SARM1 retains its ability to function as an executioner of axon degeneration. These mice are therefore a useful resource for further studies relating to SARM1 in the brain, in macrophages and in other tissues, and will allow biochemical studies to proceed involving endogenous SARM1.

Overall, this work has revealed that SARM1 is expressed in mouse macrophages but does not have a broad role in transcriptional regulation in these cells, and has provided important new animal models to further explore SARM1 function.

## METHODS

### Mice

Mice with a targeted gene deletion of exons 3-6 of *Sarm1, Sarm1^tm1Aidi^* herein referred to as B6 congenic *Sarm1*^-/-^, were generated on a C57BL/6J background from 129 ES cells and were previously described [10]. CRISPR knockout mice and epitope-tagged mice were generated as described below. Mice were housed in individually ventilated cages (IVC, Tecniplast) in a specific pathogen-free facility on a 12 h light/dark cycle with access to food and water *ad libitum* at the Trinity Biomedical Sciences Institute (TBSI), Trinity College Dublin. B6 congenic *Sarm1*^-/-^ were compared to wild-type (WT) C57BL/6J mice bred and housed at TBSI. CRISPR generated strains were compared to wild-type littermates from heterozygous breeding. Both male and female mice were used for experiments. Procedures were conducted under licenses from the Health Products Regulatory Authority (Ref: AE19136/P083) and with the approval of TCD Animal Research Ethics Committee.

### Generation of CRISPR knockout and epitope-tagged mice

#### Microinjection

Female C57BL/6J mice were superovulated with 5IU PMSG and 5IU hCG *i.p*., mated to C57BL/6J males and the recovered zygotes microinjected with pre-assembled Cas9 ribonucleoparticles (RNP) according to standard protocols [47] before being surgically transferred into the oviducts of CD1 pseudo pregnant females. *Sarm1* knockouts were made by targeting exon 1, 96 nucleotides downstream of the ATG start codon, using gRNA against the sequence GCCACCACTCCGATCCGGTC**CGG** (Figure 3A). Of the 137 zygotes microinjected, 11 offspring were recovered, screened by PCR and selected mutants characterized by Sanger sequencing (MWG eurofins). Knockout mutants used in this study had a 2 bp (*Sarm1^em1.1Tftc^*), 34 bp (*Sarm1^em1.2Tftc^*) and 5 bp (*Sarm1^em1.3Tftc^*) deletions, respectively. Epitope tagging of SARM1 involved inserting a triple FLAG, double Streptag II sequence at the C-terminus of the gene. The gRNA recognition sequence in exon 9, CACAGTAGGGCACGGGAAC**TGG**, was targeted and a repair construct used to remove the stop codon and introduce the tag-sequences (Figure 6A). Two base changes (cc > tt) were introduced at the original gRNA target site to remove the PAM sequence and prohibit recutting by Cas9. Pro-nuclear microinjection yielded 13 offspring, with 1 mutant male (*Sarm1^em2(FLAG-Strep)Tftc^*) identified by sequencing. *Sarm1^em2(FLAG-Strep)Tftc^* mice are herein referred to as *Sarm1^Flag^* mice.

#### Genotyping

Knockout CRISPR offspring were genotyped from ear DNA samples by PCR (GoTaq® polymerase) using the primer pair F1: 5’ - CATGGTCCTGACGCTGCTC - 3’, R1: 5’ - CGCCTTGCACCTCAGTGC-3’ for 30 cycles (30 s at 94 °C, 30 s at 61 °C, 30 s at 72 °C) to generate a wild-type band of 191 bp and a mutant specific band; *Sarm1^em1.1Tftc^* (189 bp), *Sarm1^em1.2Tftc^* (157 bp) and *Sarm1^em1.3Tftc^* (186 bp). All deletions were resolved from the wild-type allele by BsaWI restriction enzyme digest, which is specific to the 191 bp wild-type amplicon (Figure 3C). Genotyping of the *Sarm1^em2(FLAG-Strep)Tftc^* mice was performed by PCR using the primer pair F1: 5’ - GTACCAGGAGGCCACCATCGAG - 3’ and R1: 5’ - CTCATCTAACCTGTGCCTGGCATC - 3’ for 30 cycles (30 s at 94 °C, 30 s at 61 °C, 60 s at 68 °C) generating a wild-type band of 299 bp and an epitope specific band of 494 bp. Amplicons were electrophoresed on a 2% agarose gel.

### Cell culture of BMDM

BMDMs were cultured in DMEM + GlutaMAX (GIBCO), 10% (v/v) FCS, 100 μg/ml Normocin (InvivoGen), and 1% penicillin-streptomycin (Sigma-Aldrich), and kept at 37°C with 5% CO_2_. To generate primary BMDM (pBMDM), bone marrow from the tibiae and femurs from mice was flushed using a syringe, red blood cells were lysed, and the resulting cells were cultured in complete DMEM supplemented with 20% (v/v) L929 supernatant as a source of M-CSF [23]. On day 7, cells were removed from dishes using a sterile cell scraper and seeded for experiments in complete DMEM. Immortalized BMDMs (iBMDM) were generated with J2 recombinant retrovirus carrying ν -myc and ν -raf/mil oncogenes [48]. J2 recombinant retrovirus was kindly provided by Professor Jose Bengoechea, Queen’s University Belfast, Northern Ireland. C terminal Flag-tagged murine SARM1 was cloned into *Sarm1*^-/-^ iBMDM as previously described [25].

### Cell culture of primary neurons

Primary mouse neurons were dissected from P1-P3 neonate brains and cultured as described [49]. The cortex was dissected, trypsinized and resuspended into the Neurobasal A medium (NBAM, GIBCO) containing B27 supplement (GIBCO), 10 μM Cytosine β-D-arabinofuranoside (Ara-C, Sigma) and 0.5 mM GlutaMax supplement (GIBCO). Cells were plated in wells coated overnight with 0.1 mg/ml Poly-D-lysine (Sigma). At DIV2 (Days *in vitro*), the medium was replaced with NBAM containing B27 and one-half of the media was replaced at DIV5. Cells were used for the assay at DIV7.

### Cell treatments with stimulants

Lipopolysaccharide (LPS) serotype EH(100)Ra (ENZO) was used at 100ng/ml to stimulate TLR4 in BMDM. CL075 (InvivoGen) was used at 5 μg/ml to stimulate TLR7 in BMDM. MPLA (InvivoGen) was used at 1 μg/ml to stimulate TLR4 in BMDM.

### ELISA

Quantification of secreted CCL5, TNF, and IL-6 from cell supernatants was measured by ELISA (R&D, DuoSet ELISA kits) following the manufacturer’s instructions.

### RNA analysis by quantitative RT-PCR

For mice tissues, brain, brainstem, liver, spleen and kidney were dissected and kept in RNAlater (Invitrogen) until isolation. Total RNA was extracted from the tissues in TRIzol reagent (Ambion) according to the manufacturer’s instructions. RNA was quantified using a Nanodrop 2000 (Thermofisher). Extracted RNA was treated with DNase I (Promega) and cDNA was generated as described below. For cells in culture, cells were seeded at 5 × 10^5^ cells/ml in 24-well plates, allowed to adhere overnight, and stimulated as indicated the next day. Total RNA was extracted using the High Pure RNA Isolation Kit (Roche) and reversed transcribed with random hexamers (IDT) using Moloney murine leukemia virus (M-MLV) reverse transcriptase (Promega) according to the manufacturers’ instructions. The resulting cDNA was analyzed by quantitative RT-PCR using the PowerUp SYBR Green Master Mix (Applied Biosystems) and gene-specific primer pairs (sequences are listed in Table S1). Relative mRNA expression was calculated using the comparative C_T_ method, normalising the gene of interest to the housekeeping gene β -actin, and comparing it to an untreated wild-type sample.

### RNA sequencing

Primary BMDM were generated from femurs and tibiae of four 8-week old female WT and four *Sarm1*^-/-^ mice. Cells were stimulated for 3 hours with 5 μg/ml CL075 or left unstimulated. Total RNA was isolated prepared from samples as described above and stored at −80°C. Bioanalysis was performed using Agilent RNA 6000 Nano Kit and Agilent 2100 bioanalyzer according to manufacturer’s instructions. Library preparation and sequencing were performed by Macrogen Inc. RNA libraries were prepared using TruSeq Stranded Total RNA with Ribo-zero (Illumina). Sequencing was carried out on the Illumina NovaSeq 6000 platform, in a 100bp paired-ends read format, with a sequencing coverage of 40 million reads. Sequencing data was received from Macrogen as Fastq files. Two different forms of alignment were carried out. Hisat2 [50] was used to map reads directly to the GRCm38.p6 assembly of the C57BL/6 mouse genome, whereas kallisto [51] was used to map reads against the repeat-masked transcriptome, which were subsequently transposed onto the genome.

#### Differential gene expression analysis

Following mapping using kallisto, transcripts which were too short to have been captured at the library preparation step (<300bp in length) were filtered out of the analysis. Following this, transcripts which were very lowly expressed, having a transcript per million (TPM) count of less than one, were filtered out to remove background noise. DESeq2 [52] (version 1.22.2) was used for differential gene expression analysis of RNA sequencing data. As *Sarm1*^-/-^ replicates exhibited a lot of variability, these samples were split into two pairs with similar expression profiles (SKO1/SKO2 and SKO3/SKO5) then compared to WT. Unstimulated *Sarm1*^-/-^ and WT pBMDM were compared, and genes were deemed differentially expressed if the DESeq2 base mean was greater than 100, the fold change exceeded 1.6 fold, and the adjusted p-value was less than 0.05. Similarly, CL075-stimulated *Sarm1*^-/-^ and WT pBMDM were compared, and genes were deemed differentially expressed if the DESeq2 base mean was greater than 100, and the fold change exceeded 1.6 fold. The p-value cut off for two replicates (SKO3 and SKO5) was relaxed to p≤0.1 to include *Ccl5*, which was the positive control. A heatmap was then generated using the ComplexHeatmap [53] package in R, showing genes which were differentially expressed either before or after stimulation in both SKO1/2 and SKO3/5 relative to WT, grouped according to chromosomal location.

#### Analysis of density of SNPs and indels relative to the reference genome

The samtools mpileup [54] tool was used to identify genetic variants between a representative unstimulated *Sarm1*^-/-^ sample sequence (mapped using Hisat2) and the mm10 mouse reference genome. Genetics variants (both SNPs and INDELS) were filtered to only include: 1. Those with high quality score (QUAL > 100) and 2. Those with high sequencing depth (DP > 100). The mouse reference genome was divided into sliding windows of 10kb using bedtools [55] makewindows. Genetic variants were counted within each of these windows using bedtools intersect. The count data was visualized in R with the IdeoViz package.

#### Visualizing gene coverage, SNPs, and indels using Integrative Genomics Viewer (IGV)

The integrative genomics viewer [56] (IGV) from the Broad Institute was used to visualize reads which were mapped to the genome using Hisat2. BAM files of these reads were loaded to IGV so that sequence variation from the reference sequence (denoted by a coloured bar) and gene coverage could be examined at a given locus.

### Cell Fractionation

Cells were seeded at 5 × 10^5^ cells/ml in 10 cm culture dishes and allowed to adhere overnight. The next day, cells were stimulated as indicated. Following stimulation, supernatants were removed and discarded, and cells were washed twice with ice-cold phosphate buffered saline (PBS) on ice. Cells were scraped in 5 ml ice-cold PBS then pelleted by centrifugation at 1,000g for 5 minutes at 4 °C. Cells were resuspended in 200 μl cytosolic lysis buffer (10 mM Tris-HCl pH 7.4, 10 mM NaCl, 3 mM MgCl_2_, 10 mM EDTA, 1% Triton X-100) supplemented with 1 mM sodium orthovanadate, 10 μl/ml aprotinin (Sigma), and 1 mM PMSF (protease inhibitors). The cell suspension was incubated for 30 minutes on ice to lyse the plasma membrane, then centrifuged for 30 min at 2,000 rpm at 4 °C. The supernatant, containing an impure cytoplasmic fraction, was removed and stored at −80 °C. The pellet, containing a pure nuclear fraction was washed twice with 1 ml of unsupplemented cytoplasmic lysis buffer, then resuspended in 200 μl nuclear lysis buffer (30 mM HEPES pH 7.9, 36% glycerol, 600 mM NaCl, 2 mM MgCl_2_, 30 mM EDTA). During a 30 min incubation on ice, samples were vigorously vortexed three times. The samples were then centrifuged at 13,200 rpm. Supernatants containing nuclear proteins was removed and stored at −80 °C.

### Immunoblotting

The antibodies used were as follows. Primary antibodies specific for SARM1 (Chicken polyclonal IgY, established in our laboratory), FLAG (Sigma, F1804), Vinculin (CST, #4650), β-actin (Sigma-Aldrich, A5316), GAPDH (Cell Signaling Technology, #97166), H3 (Upstate, 05-499). Anti SARM1 Antibody was generated by immunizing chicken with the TIR domain of human SARM1 recombinant protein tagged with 6xHis-tag [4]. The recombinant protein was kindly provided by Prof. Bostjan Kobe (University of Queensland, Brisbane, Australia). We also used IRDye anti-chicken IgY, anti-mouse IgG and anti-rabbit IgG (Li-cor) as secondary antibodies for immunoblotting.

Mice tissues, brain and brainstems were dissected and homogenized in ice-cold lysis buffer (50 mM Tris-HCl, pH8.0, 150 mM NaCl, 5 mM EDTA, 1 % Triton X-100, and cOmplete™ mini Protease Inhibitor Cocktail (Roche)). Following incubation on ice for 30 min, samples were centrifuged for 20 min at 15,000 rpm at 4°C. The supernatant (lysates) were transferred to a fresh tube, mixed with Laemmli sample buffer and boiled for immunoblotting.

BMDM were seeded at 5 × 10^5^ cells/ml and allowed to adhere overnight. BMDM were washed with PBS three times, then lysed with RIPA buffer (50 mM Tris-HCl, pH 8, 150 mM NaCl, 1% Triton X-100, 0.1% SDS, 0.5% sodium deoxycholate, 5 mM EDTA) supplemented with 1mM sodium orthovanadate, 10 μl/ml aprotinin, and 1 mM PMSF. Prior to preparing samples for Western Blot, protein concentrations were measured by bicinchoninic acid (BCA) assay (Pierce) according to the manufacturer’s instructions. Lysates were mixed with Laemmli sample buffer and boiled for 5 minutes at 99 °C.

SDS-PAGE was carried out, and proteins were transferred to a nitrocellulose membrane using the semidry transfer method. Membranes were blocked using a 5% (w/v) solution of skimmed milk powder in PBS-Tween (0.1% Tween, v/v), then incubated with antibodies as indicated. Membranes were read on the Li-Cor Odyssey, and images were adjusted using Image Studio Lite ver. 4.

### Immunoprecipitation

WT and *Sarm1^flag^* iBMDM were seeded at 5 × 10^5^ cells/ml in 15 cm dishes and allowed to adhere overnight. The following day, supernatants were removed and cells were washed three times with ice-cold PBS. Cells were then lysed using IP lysis buffer (50 mM Tris-HCl, pH 7.4, with 150 mM NaCl, 1 mM EDTA, and 1% Triton X-100) supplemented with 1 mM sodium orthovanadate, 10 μl/ml aprotinin, and 1 mM PMSF. Immunoprecipitation was carried out using anti-FLAG M2 Magnetic Beads (Sigma, M8823) according to the manufacturer’s instructions. Immunoprecipitated proteins were eluted from the beads by incubation with glycine-HCl (0.1 M, pH 3), then neutralized with 1 M Tris-HCl. IP lysates and input controls were subject to SDS-PAGE as described above.

### Axon degeneration and measuring length of the neurite

Neurons (DIV7) were treated with 1 nM vincristine sulfate salt (Sigma) or vehicle (DMSO), and then were set into Incucyte Live-Cell Analysis Systems (Sartorius). The first scan was made at 1h after the treatment, the following scans were made every 6h after the treatment. The neurite outgrowth was quantified using the semi-automatic tracing tool NeuronJ plugin [57] from the ImageJ package Fiji [58].

### Statistical analysis

All data were analyzed with GraphPad Prism 9. Data are presented as mean ± SEM; *p < 0.05, **p < 0.01, ***p < 0.001, ****p < 0.0001. Specific statistical tests are described in detail in figure legends.

## ACKNOWLEDGEMENTS

We thank Katharine A. Shanahan for technical assistance with mice and genotyping. Imaging works were performed at the TBSI Microscopy and Imaging Centre, with technical assistance from Gavin McManus. We thank members of the Bowie lab for helpful discussions. This work was funded by Science Foundation Ireland (16/IA/4376 to A.G.B. and 05/IN3/B761 to V.P.K.).

## AUTHOR CONTRIBUTIONS

Conceptualization, A.G.B.; Methodology, C.G.D., R.S., F.R., K.H., C.F., V.P.K.; Investigation, C.G.D., R.S., M.C., F.R., K.H.; Resources, C.F., V.P.K.; Writing – Original Draft, C.G.D, R.S., A.G.B.; Supervision, M.C., K.H., V.P.K., A.G.B.; Funding Acquisition, A.G.B.

## SUPPLEMENTAL INFORMATION

Supplemental information includes 1 table, 7 figures and 2 movies.

## DATA AVAILABILITY

All data generated or analyzed during this study are included within this article (and its supplementary information files). RNA sequencing data is available as GEO data set GSE182091 at https://www.ncbi.nlm.nih.gov/geo/query/acc.cgi?acc=GSE182091

