## Supplemental Table and Figures for "Next generation SARM1 knockout and epitope tagged CRISPR-Cas9-generated isogenic mice reveal that SARM1 does not participate in regulating nuclear transcription, despite confirmation of protein expression in macrophages"

**Supplementary Table 1:** List of primers used for quantitative RT-PCR and genotyping

| **Target** | **Sequence (5'---3')** |
|---|---|
| *Tnf (forward)* | TCC CCA AAG GGA TGA GAA GTT |
| *Tnf (reverse)* | GTT TGC TAC GAC GTG GGC TAC |
| *Ccl5 (forward)* | CTC ACC ATA TGG CTC GGA CA |
| *Ccl5 (reverse)* | ACA AAC ACG ACT GCA AGA TTG G |
| *Il6* (forward) | AAG AGT TGT GCA ATG GCA ATT CTG |
| *Il6* (reverse) | ATA GGC AAA TTT CCT GAT TAT ATC CAG T |
| β-*actin* (forward) | TCC AGC CTT CCT TCT TGG GT |
| β-*actin* (reverse) | GCA CTG TGT TGG CAT AGA GGT |
| *Ccl6* (forward) | CTT TAT CCT TGT GGC TGT CC |
| *Ccl6* (reverse) | TGA ATT ATT GGA GGG TTA TAG CG |
| *Ccl9* (forward) | TAA CTC ACG GAT TCA GTG TTC |
| *Ccl9* (reverse) | CCA ATC TTT CAA TGC ATC TCT G |
| *Rps29* (forward) | AGC TCT ACT GGA GTC ACC |
| *Rps29* (reverse) | TTC AGC CCG TAT TTG CG |
| *Rpl38* (forward) | GGA GAT CAA GGA CTT TCT GC |
| *Rpl38* (reverse) | GTG ATA ACC AGG GTG TAA AGG |
| *Uqcrh* (forward) | GTG TCT GAT GGA GTG AGT TC |
| *Uqcrh* (reverse) | CCT GAT TCC CAG TGA CAA G |
| *Atp5k* (forward) | CAT CGG CAT GGC ATA CG |
| *Atp5k* (reverse) | GCT GTC ATC TTG AGC TTC C |
| *Sarm1* (forward) | GCT CAG TGC ATA GGA GCA TTC |
| *Sarm1* (reverse) | AGT AAG AAA CCA GGC GTT TCA G |
| *Acap1* (forward) | GGC ATT GTC AGA TCC AAA TC |
| *Acap1* (reverse) | CAT CAG CCA TGG TAG GAA G |
| *Ndufb3* (forward) | TGT CGT AAG AAA CTA GAG GAA AC |
| *Ndufb3* (reverse) | CAT GTC CAT GTC CAG CAG |
| *Tfrc* (forward) | CTC GCT TAT ATT GGG CAG AC |
| *Tfrc* (reverse) | CTC ACG AGG AGT GTA TGT ATT C |
| *Sarm1*$^{em1.1Tftc}$, *Sarm1*$^{em1.2Tftc}$ and *Sarm1*$^{em1.3Tftc}$ genotyping (forward) | CAT GGT CCT GAC GCT GCT C |
| *Sarm1*$^{em1.1Tftc}$, *Sarm1*$^{em1.2Tftc}$ and *Sarm1*$^{em1.3Tftc}$ genotyping (reverse) | CGC CTT GCA CCT CAG TGC |
| *Sarm1*$^{Flag}$ genotyping (forward) | GTA CCA GGA GGC CAC CAT CGA G |
| *Sarm1*$^{Flag}$ genotyping (reverse) | CTC ATC TAA CCT GTG CCT GGC ATC |

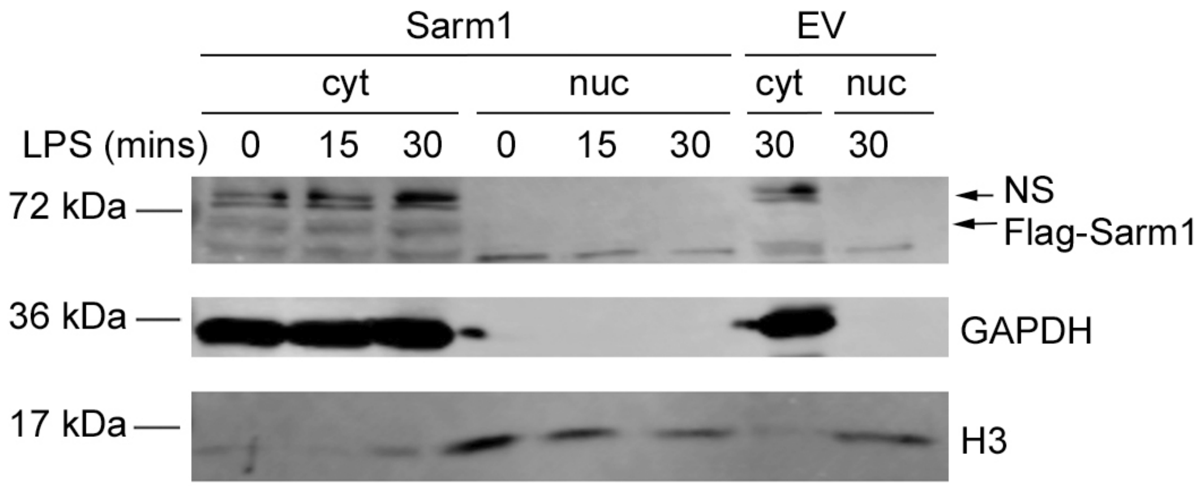


**Figure S1. SARM1 is expressed in the cytoplasm and not the nucleus.** Related to Figure 1.
*Sarm1*$^{-/-}$ iBMDM stably expressing empty vector (EV) or flag-tagged SARM1 were stimulated with 100 ng/ml LPS as indicated, then a nuclear fraction (nuc) and cytoplasmic (cyt) fraction were isolated by fractionation and subjected to SDS-PAGE. Immunoblotting was performed for SARM1 using a specific antibody to flag, with H3 as a nuclear marker, and GAPDH as a cytoplasmic marker. NS indicates a non-specific band. Blot is representative of three independent experiments.

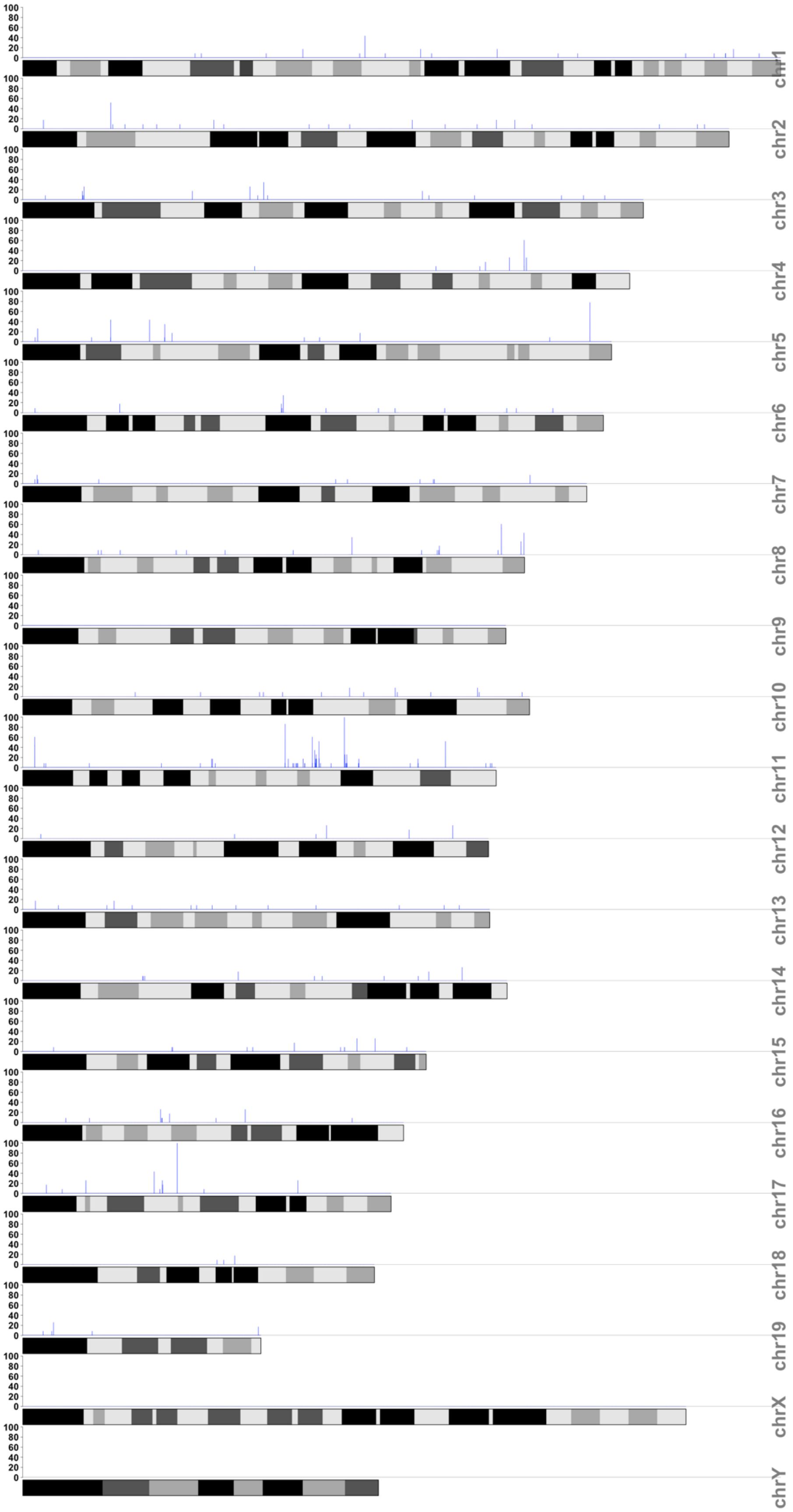


**Figure S2. Density of SNPs and indels on the full mouse chromosome set from *Sarm1*$^{-/-}$ mice.** Related to Figure 2. Graph showing density of SNPs and indels per 10 kb section of each chromosome in *Sarm1*$^{-/-}$ BMDM relative to the C57BL/6 reference sequence.

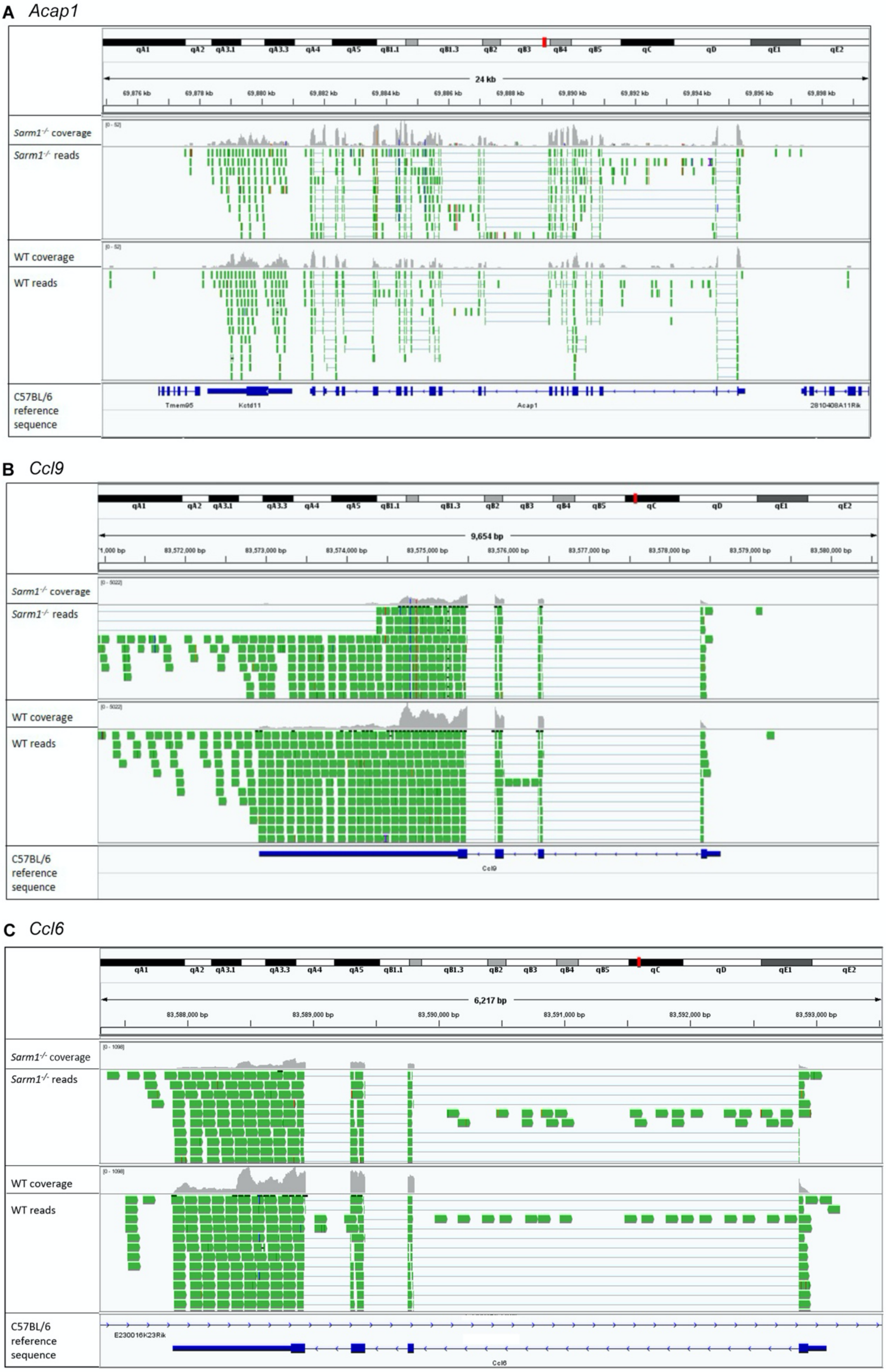


**Figure S3. IGV tracks of selected genes from chromosome 11 reveal sequence variation between *Sarm1*$^{-/-}$ and WT mice.** Related to Figure 2. Aligned reads at the *Acap1* (A), *Ccl9* (B), and *Ccl6* (C) loci in unstimulated WT and *Sarm1*$^{-/-}$ BMDM were visualized using IGV. Images show coverage of the genes in grey, with variations from the C57BL/6 reference sequence denoted by coloured vertical bars.

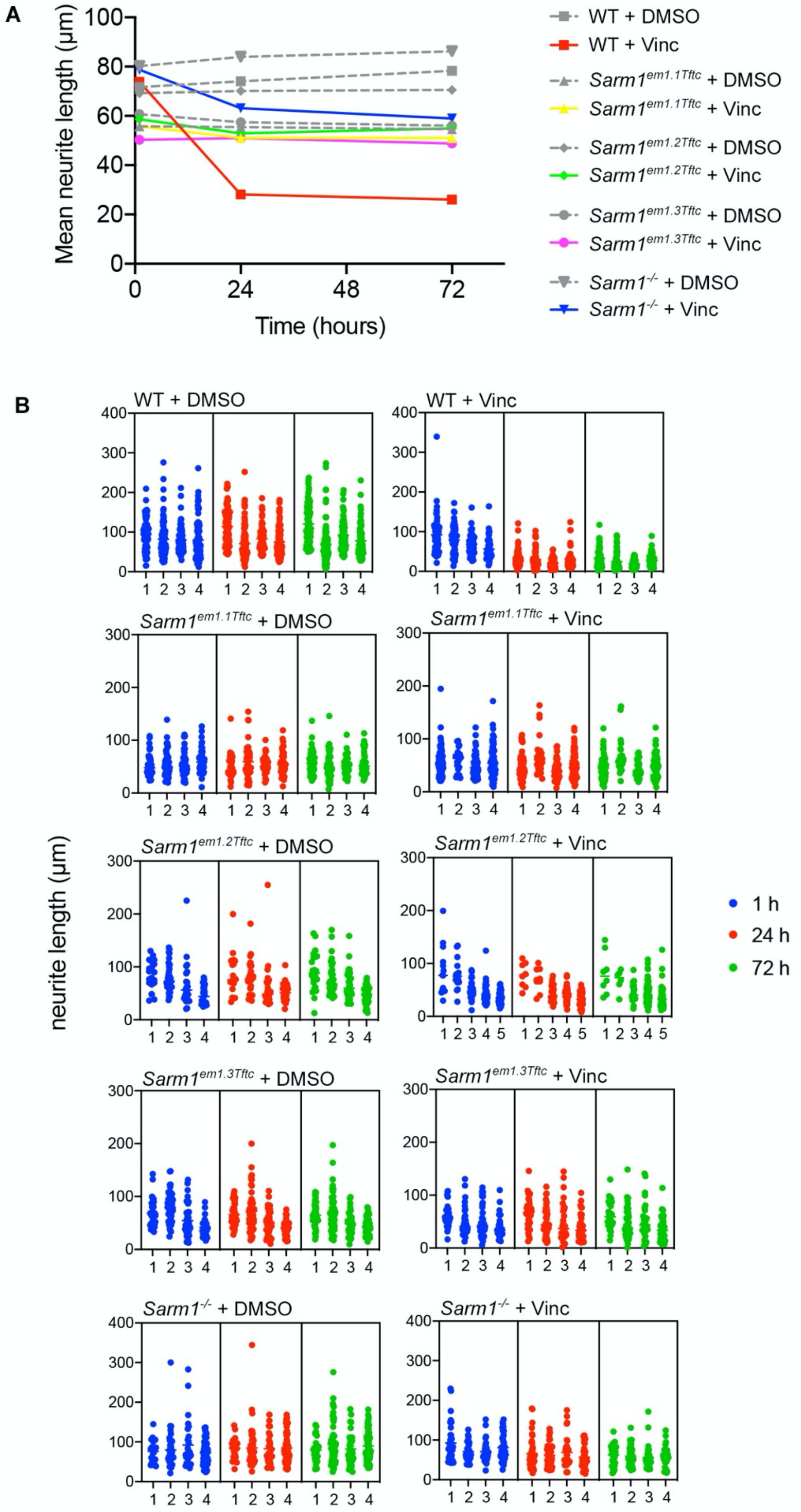


**Figure S4. Analysis of neurite length following vincristine treatment in neurons from *Sarm1* knockout mice.** Related to Figure 3. (A) Graph of raw neurite length over time for different mice and treatments shown in Figure 3F. All data are mean per head ± SEM (n= 4~5). (B) Neurite length data used for analysis in Figure 3F. Each number on x-axis indicates an individual head and each dot shows neurite length per neuron.

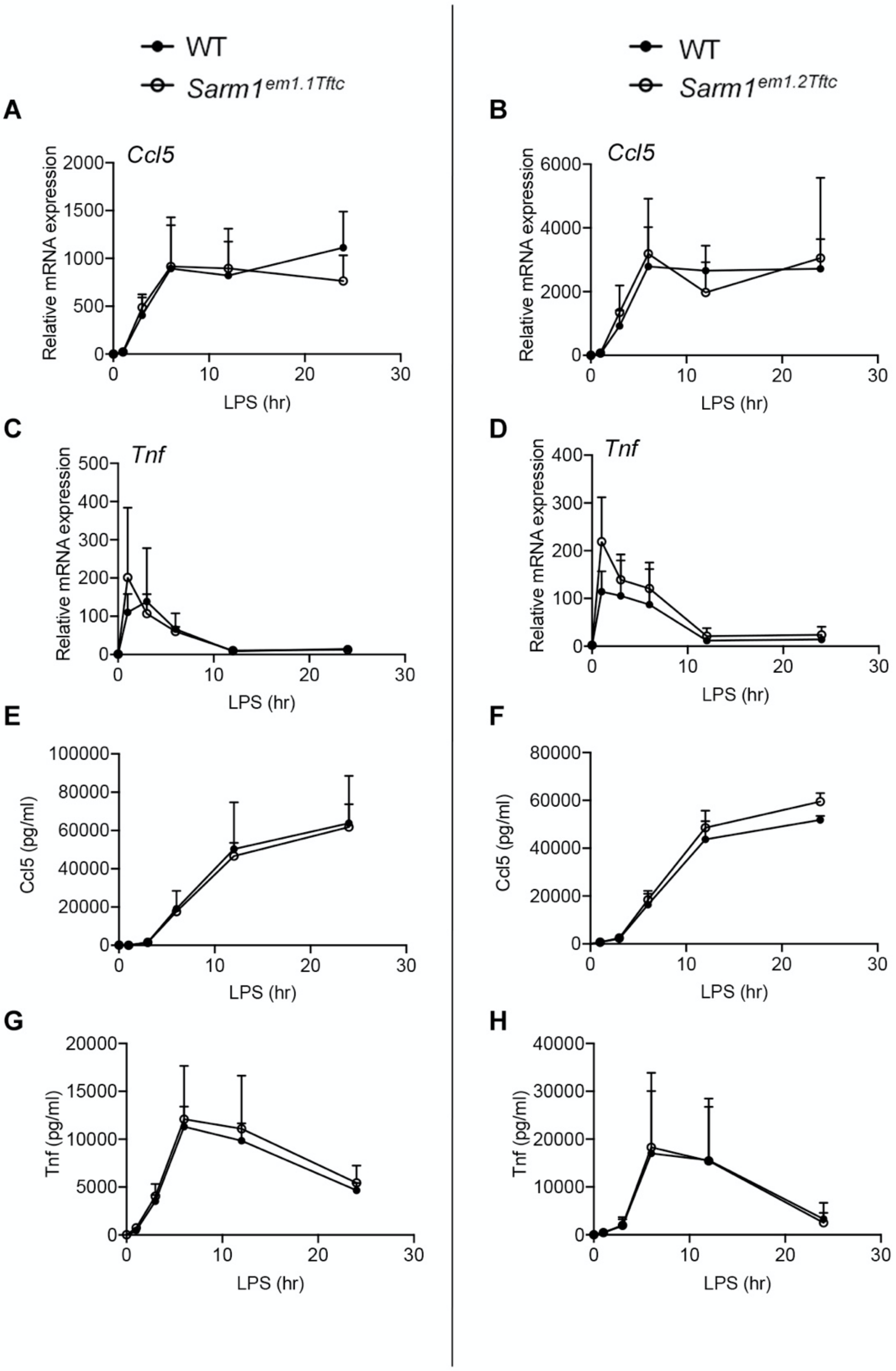


**Figure S5. Normal induction of *Ccl5* and *Tnf* in macrophages from $Sarm1^{em1.1Tftc}$ and $Sarm1^{em1.2Tftc}$ mice compared to WT littermates.** Related to Figure 4. WT and $Sarm1^{em1.1Tftc}$ or $Sarm1^{em1.2Tftc}$ pBMDM were stimulated with 100 ng/ml LPS for the indicated times, or medium as a control. Expression of *Ccl5* (A, B) and *Tnf* (C, D) mRNA were assayed by qRT-PCR, normalized to the housekeeping gene β-actin, and are presented relative to the untreated WT control. Supernatants were assayed for CCL5 (E, F) and TNF (G, H) protein by ELISA. Data are mean ±SEM (n=2). No significant differences determined by multiple Mann-Whitney tests; Holm-Šídák multiple comparisons.

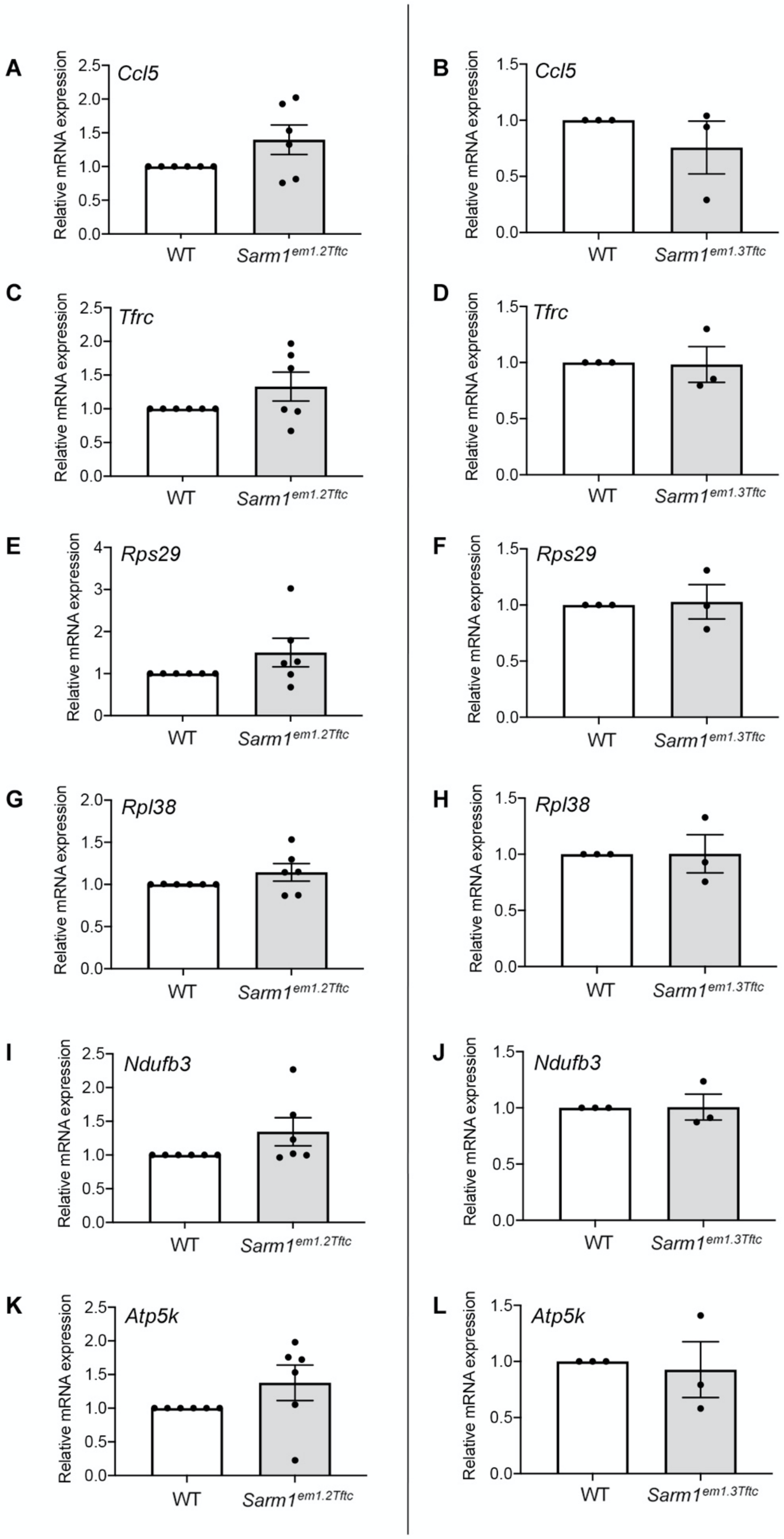


**Figure S6. Basal transcription of select brainstem genes is unaltered in *Sarm1*$^{em1.2Tftc}$ and *Sarm1*$^{em1.3Tftc}$ mice compared to WT littermates.** Related to Figure 5. Expression of *Ccl5* (A, B), *Tfrc* (C, D), *Rps29* (E, F), *Rpl38* (G, H), *Ndufb3* (I, J), and *Atp5k* (K, L) in the brainstem of *Sarm1*$^{em1.2Tftc}$ and *Sarm1*$^{em1.3Tftc}$ mice respectively were measured by qRT-PCR, normalized to β-actin, and are presented relative to a littermate WT. Data are mean ±SEM (n=6, *Sarm1*$^{em1.2Tftc}$ no significant differences determined by multiple Wilcoxon tests; Holm-Šídák multiple comparisons, n=3 *Sarm1*$^{em1.3Tftc}$).

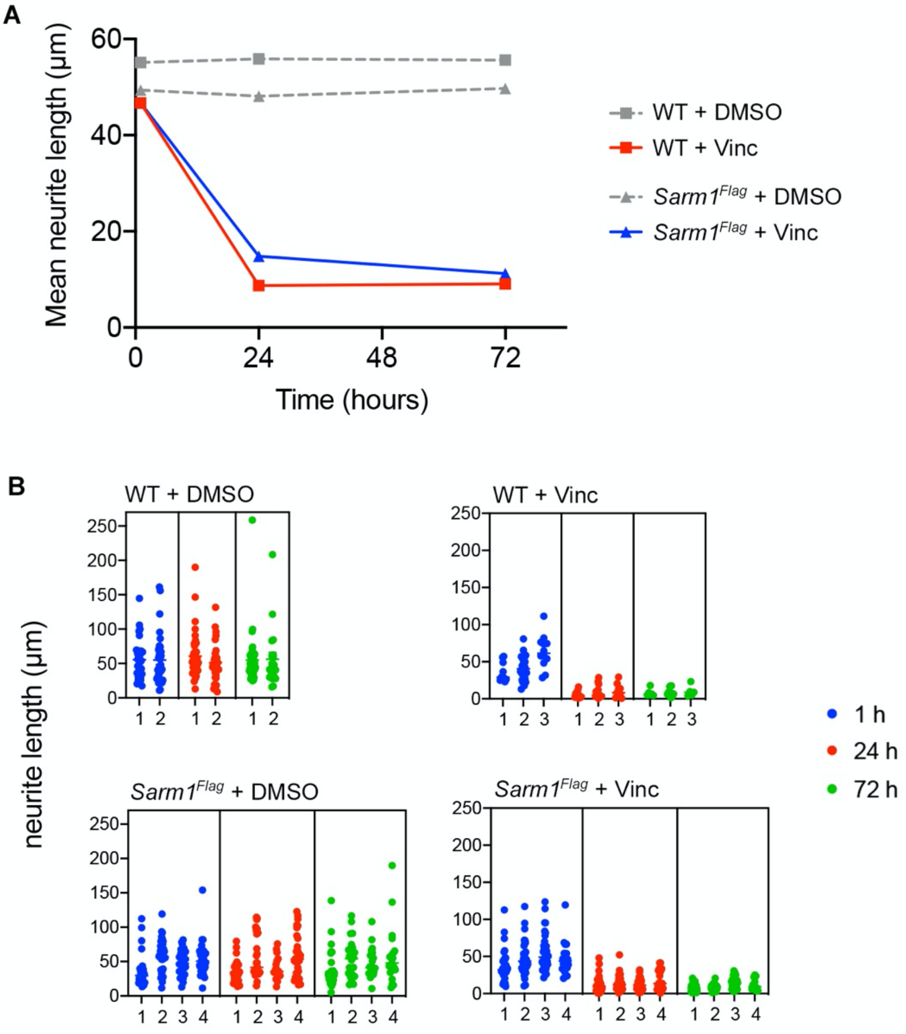


**Figure S7. Analysis of neurite length following vincristine treatment in neurons from *Sarm1$^{Flag}$* mice.** Related to Figure 7. (A) Graph of raw neurite length over time for different mice and treatments shown in Figure 7E. All data are mean per head ± SEM (n= 2~4). (B) Neurite length data used for analysis in Figure 7E. Each number on x-axis indicates an individual head and each dot shows neurite length per neuron.

**Supplemental movies S1 & S2. The effect of vincristine treatment on WT neurons.** Related to Figure 3. Representative images of neurons after treatment of DMSO (S1) or vincristine (S2). Images were scanned from 1h after the treatment, which is indicated as 0h in the movie, and following every 6h from 6h after the treatment (shown as 5h) until 72h (shown as 71h).
